# Omission-responsive neurons encode negative prediction error and probability in the auditory cortex

**DOI:** 10.1101/2024.09.09.612166

**Authors:** Amit Yaron, Tomoyo Shiramatsu-Isoguchi, Felix B. Kern, Kenichi Ohki, Hirokazu Takahashi, Zenas C. Chao

## Abstract

Predictive coding posits the brain predicts incoming sensory information and signals prediction errors when actual input differs from expectations. Positive prediction errors occur when input exceeds predictions, while negative prediction errors arise when input falls short. Specific neurons are theorized to encode negative prediction errors, linked to responses to omitted expected inputs. However, the information encoded in omission responses remains unclear. We recorded single-unit activity in rat auditory cortex during an omission paradigm with varying tone probabilities. We identified neurons selectively responding to omissions, with responses increasing with evidence accumulation and correlating with tone predictability—key characteristics of negative prediction-error neurons. Interestingly, these neurons showed selective omission responses but broad tone responses, revealing an asymmetry in error signaling. We propose a circuit model with laterally interconnected prediction-error neurons reproducing this asymmetry. Our model demonstrates that lateral connections enhance precision and efficiency of prediction encoding, supported by the free energy principle.

## Introduction

Predictive coding, a theoretical framework that explains how the brain processes sensory information, posits that the brain generates predictions based on prior experiences and compares them with actual sensory inputs [1–5]. The discrepancy between the predicted and actual input, known as the prediction error, is used to update the brain’s internal models. This process is thought to occur through canonical circuits, hierarchically organized neural networks where top-down predictions and bottom-up prediction errors create a feedback loop that allows the brain to refine its models.

Recent work suggests that positive prediction errors, occurring when the actual input exceeds the predicted input, and negative prediction errors, occurring when the actual input is less than the predicted input, are calculated in separate populations of neurons within these canonical circuits [6–9]. While extensive research has investigated positive prediction errors using the oddball paradigm and its variations [10–18] these studies face significant limitations. Specifically, oddball responses consist of more than just positive prediction errors. First, they also contain sensory signals, which are often hard to disentangle. Additionally, a deviant stimulus also represents the omission of the standard stimulus, thereby generating a negative prediction error, which further complicates the interpretation. In contrast, responses to omitted expected stimuli do not contain sensory input and could directly represent negative error signals, potentially offering a crucial step in elucidating the computation of prediction errors. Omission responses have been observed in human studies [19–24] and more recently in single neurons within the visual and auditory cortices [7,25–27]. However, omission responses could arise from processes other than predictive coding and may not necessarily represent negative prediction errors. Therefore, it is crucial to characterize the information encoded in these responses to confirm their functional role and investigate how they might arise from the interactions between prediction and sensory signals, an area that remains largely unexplored.

To address this, we investigated how neurons in the rat auditory cortex respond to the omission of expected auditory stimuli while manipulating their predictability. We use high-resolution extracellular recordings to simultaneously record activity from a large population of neurons across multiple cortical layers, while presenting tone sequences where the probability of tone presentations was systematically varied. We identified a subset of auditory neurons that exhibit two key characteristics of negative prediction-error encoding. First, these neurons exhibit omission responses that build up from trial to trial, reflecting predictions established by the accumulation of evidence through repeated exposure to the stimuli. Second, the neurons displayed omission responses whose intensity was directly correlated with the statistical likelihood of specific omitted tones, demonstrating a quantitative representation of the prediction error. These neurons, which we named “Probability Encoding Omission Neurons” (PEONs), were primarily found in the granular and supragranular layers of the auditory cortex and were evenly distributed across different auditory fields including primary auditory cortex (A1), ventral auditory field (VAF), and anterior auditory field (AAF).

Interestingly, while PEONs showed selective responses to omissions, they responded broadly to tones, highlighting an asymmetry between the processing of top-down predictions and bottom-up sensory signals. To explain these dynamics, we propose a novel circuit model that includes separate negative and positive error calculations within narrowly tuned auditory streams, with prediction error neurons laterally interconnected between adjacent streams. This architecture not only accounts for the observed asymmetry and probability encoding in sensory processing but also demonstrates how lateral connections can enhance the precision and efficiency of prediction-encoding, a concept supported by the free energy principle [1,3]. Our findings provide empirical evidence for the existence of negative prediction error neurons in the auditory cortex and offer a comprehensive account of how these errors are computed and integrated within cortical microcircuits.

## Results

### Neuronal Responses During Auditory Omission Paradigm

In this study, we focused on the right auditory cortex of Urethane-anesthetized rats, utilizing extracellular recordings to monitor responses to auditory stimuli. Across eighteen electrode penetrations involving 10 rats, we successfully captured the activity of 1,489 single units. For each rat, we first used a surface microelectrode array (NeuroNexus, Ann Arbor, MI, USA) (Figure 1A) to map the various parts of the auditory cortex (Figure 1B)[28–30]. We then used a Neuropixels electrode array to record different layers of the chosen area for each penetration (Figure 1C).

**Figure 1.**
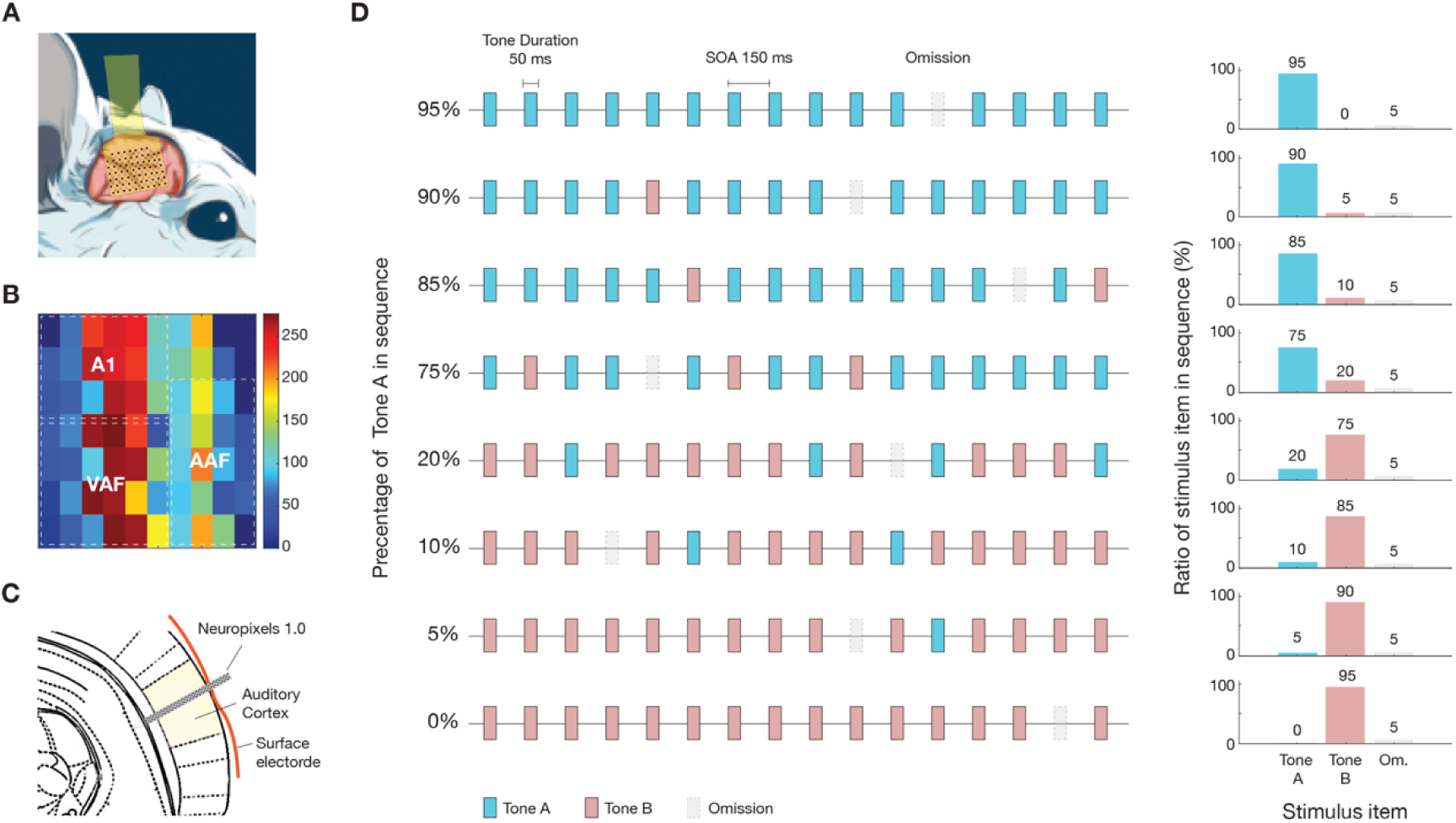
Neuronal recording and experimental paradigm. (**A**) Surface microelectrode array used to map the auditory cortex in urethane-anesthetized rats. (**B**) Representative auditory-evoked potentials in the cortex elicited by click stimuli. The color bar represents P1 amplitude (see Methods). A1, AAF, and VAF represents the primary auditory cortex, anterior auditory field, and ventral auditory field, respectively. (**C**) Neuropixels electrode array inserted through the holes on the surface array into the mapped regions to record neural activity across different cortical layers. (**D**) Experimental design featuring eight conditions with two distinct tones (A and B) and a 5% omission rate, presented in random order. Each condition comprised 1000 stimulus items, varying the probability of Tones A and B to achieve different predictability levels. Bars on the right side illustrate the tone probabilities and omission rates for each condition, ranging from 95% Tone A and 0% Tone B (exclusively Tone A) to 0% Tone A and 95% Tone B (exclusively Tone B), with a consistent 5% omission rate across all conditions.

To explore the neural response to auditory stimuli and their omission, we devised eight experimental conditions (Figure 1D). Each condition featured an oddball sequence of two tones with distinct probabilities, along with a consistent occurrence of tone omissions at a rate of 5%. To create varied prediction contents while maintaining a fixed omission rate, we utilized two tones and controlled their relative predictability. We selected two distinct pure tones (denoted as Tones A and B), one octave apart, that consistently elicited responses from most neurons for each penetration (see details in Methods). The sequences were systematically designed but presented in a random order. Each sequence comprised 1000 stimulus items, including omissions, and featured Tones A and B at varying probabilities to achieve distinct levels of predictability. The specific ratios of Tones A and B were carefully controlled as follows: 95% Tone A and 0% Tone B (exclusively Tone A), 90% and 5%, 85% and 10%, and 75% and 20%. This pattern continued, adjusting further to 20% and 75%, 10% and 85%, 5% and 90%, until reaching 0% and 95% (exclusively Tone B).

Using the responses to these sequences, we identified a substantial subset of 857 units as auditory neurons. This classification was based on comparing neuronal activity recorded just before each trial (−25 to 5 ms, where time zero represents the stimulus onset) involving either a tone or an omission, with the activity during the stimulus or expected stimulus window (5 to 120 ms). Neurons were classified as auditory if they responded significantly to at least one of the tones or their omission in one or more of the eight experimental conditions, as determined by the Benjamini-Hochberg procedure to adjust for false discoveries (see details in Methods).

### Selective Neural Encoding of Auditory Omissions

We observed diverse patterns of neural responses to omissions. Figure 2 illustrates responses from three exemplary neurons to tone omissions during two distinct tone sequences, one with 95% Tone A and 5% omissions and the other with 95% Tone B and 5% omissions. This figure highlights the varied neural reactions to the absence of expected auditory stimuli. Some neurons exhibited strong responses either tone was presented but did not respond to their omission (Figure 2A). Other neurons showed pronounced sensitivity to omissions, reacting strongly when either tone was omitted (Figure 2B). Additionally, some neurons displayed selective responses to the absence of specific tones, with the intensity and nature of these responses varying depending on which tone was omitted (Figure 2C).

**Figure 2.**
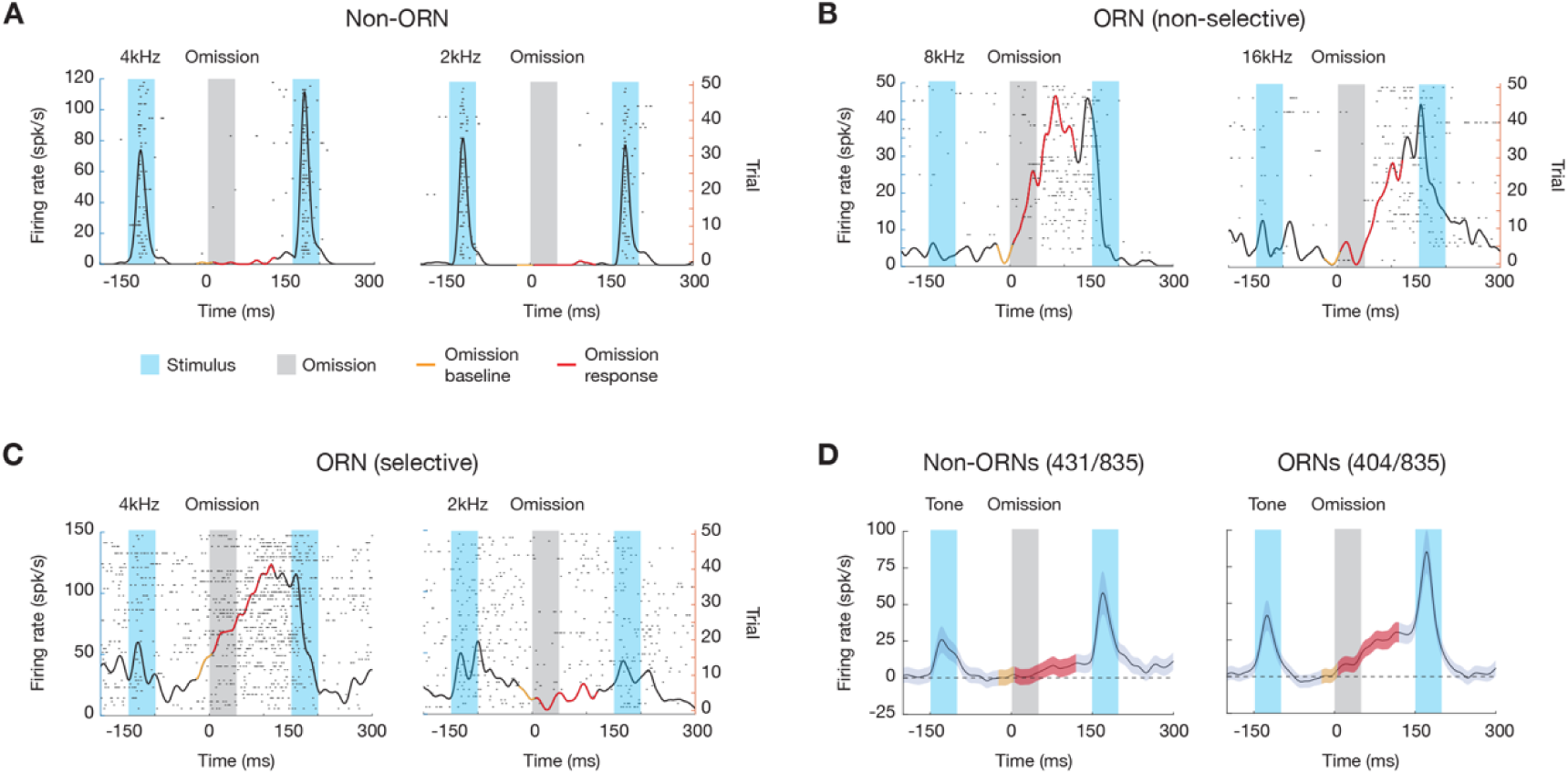
Neural responses to auditory omissions. (**A**) Non-ORN: Example of a neuron that responds to tones but not to omissions. The raster plot shows spikes (black dots) with time (ms) on the x-axis and trials on the y-axis. The peristimulus time histogram (PSTH) displays firing rate over time, with tone stimuli (blue shading) and omissions (gray shading). The black line represents the firing rate, the red line shows omission responses, and the orange line indicates the omission baseline. (**B**) ORN (non-selective): Example of a neuron that responds to omissions regardless of tone frequency. The same representation is used as in panel A. (**C**) ORN (selective): Example of a neuron that selectively responds to the omission of specific tones. The same representation is used as in panel A. (**D**) Grand-average PSTH for non-ORNs and ORNs. The left panel shows the average response of non-ORNs (431 out of 835 neurons) with significant tone responses (blue shading) but modest responses during omissions (gray shading). The right panel displays the average response of ORNs (404 out of 835 neurons) with pronounced omission responses (red shading) characterized by a slow build-up to peak, indicating stronger and more sustained responses to omissions compared to tones. The shaded areas represent standard error of the mean.

We identified neurons that demonstrated a significant response to the absence of anticipated tones in at least one of the eight experimental conditions. These neurons are referred to as “Omission Responsive Neurons” (ORNs). Remarkably, ORNs comprised a substantial fraction of the auditory neurons we identified, numbering 411 out of a total of 857 auditory neurons analyzed (approximately 48%). Figures 2D shows the grand-average peristimulus time histogram (PSTH) for non-ORNs and ORNs. For non-ORNs, there was a significant response to tones, as confirmed by a one-sided Wilcoxon signed-rank test (p= 2.3e-07, z= 5.04). The tone responses were sharp, reaching a peak rapidly. During tone omissions, we observed a modest but significant increase in activity (p= 3.7e-05, z= 3.96), indicating that the non-ORN population as a whole still exhibited responses above baseline, despite the absence of significant responses on an individual level. In contrast, ORNs demonstrated a marked and significantly stronger response to tone omissions (p= 5.3e-23, z= 9.81). Notably, this response was characterized by a slow build-up to peak. ORNs also showed a significant response to tones (p= 6.5e-07, z= 4.84). Because of the high prevalence of tone selectivity of ORNs, we defined the tone that elicited the strongest omission response from each neuron as the “Omission Preferred” (O_P_) tone, and the other tone as the “Omission Non-Preferred” (O_NP_) tone.

### Omission Responses Build Up Throughout the Trials

To confirm that ORNs encode negative prediction errors, it is essential to show that their responses to omissions increase over time as predictions are formed, reflecting a growing discrepancy between the predicted and actual outcomes. Since ORNs responded to the O_P_ tone, we analyzed their responses to its omission across sequences where the O_P_ tone was more frequent (95%, 90%, 85%, and 75%). Our analysis revealed a notable pattern: the magnitude of the ORNs’ response to these omissions progressively increased throughout the sequence, achieving a steady state or plateau after approximately the 20th omission (Figure 3A). The dynamics of the trial-by-trial omission response were fitted by a logistic growth curve, expressed as

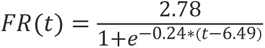

where *FR(trial)* represents the firing rate at trial *t*. The logistic function depicts a saturating increase in response magnitude. Here, an R^2^ value of 0.39 indicates the model’s moderate success in capturing the variability in responses. To demonstrate that the observed increase in omission response was not due to network activation drifting over time, we analyzed the ORNs’ responses to auditory stimuli. The results indicated a rapid decline in response intensity over successive trials (Figure 3B). This decay was fitted using an exponential decay model

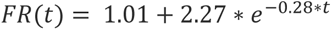

with an R^2^ value of 0.67, indicating a substantial fit to the data. As expected, this encapsulates the phenomenon of stimulus-specific adaptation or a decrease in positive prediction error, where the response to an expected stimulus diminishes over time.

**Figure 3.**
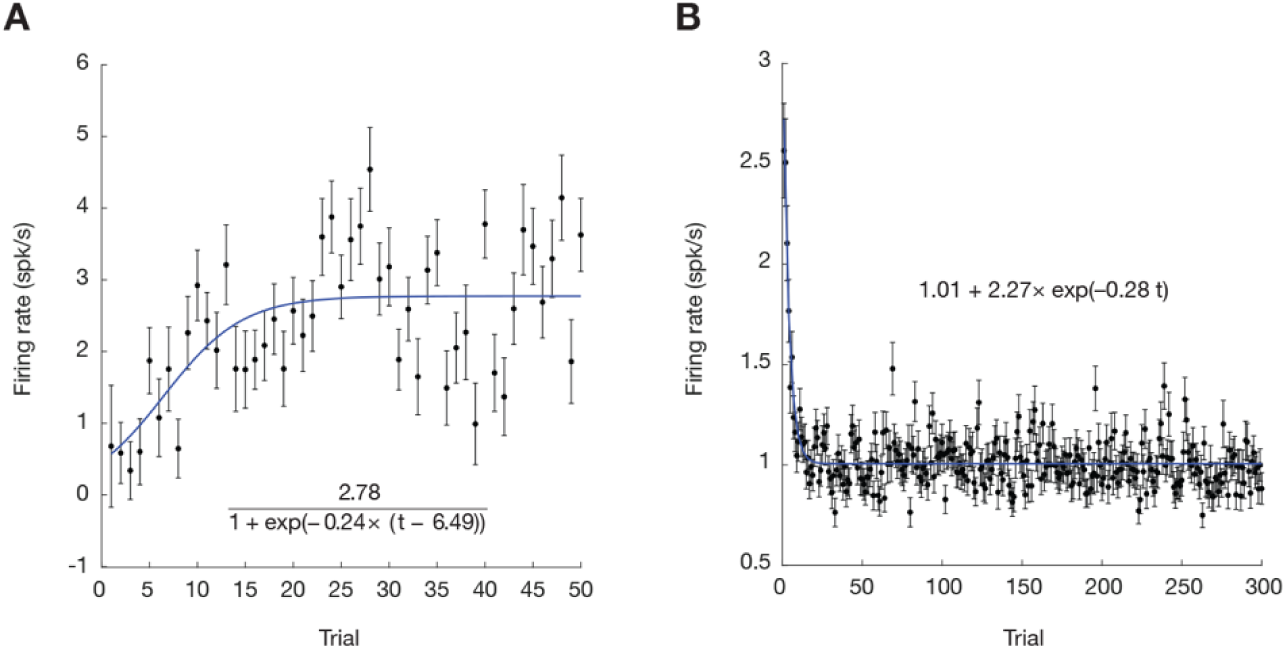
Trial-by-trial dynamics of omission responses. (**A**) Firing rate of ORNs in response to the omission of the OP tone across 50 trials. The x-axis represents the trial number, and the y-axis represents the firing rate in spikes per second. The black dots indicate the mean firing rate across all ORNs for each trial, and the error bars represent the standard error of the mean (SEM), showing variability across neurons. The blue line represents the logistic growth curve fit to the data, indicating a progressive increase in response magnitude as the neurons learn to predict the omission over time. Data were averaged across sequences where the OP tone was the standard (95%, 90%, 85%, and 75%). (**B**) Firing rate of ORNs in response to the presentation of tones across 300 trials. The x-axis represents the trial number, and the y-axis represents the firing rate in spikes per second. The black dots indicate the mean firing rate across all ORNs for each trial, and the error bars represent the standard error of the mean (SEM), showing variability across neurons. The blue line represents the exponential decay model fit to the data, indicating a rapid decline in response intensity over the initial trials as the neurons adapt to the repeated tone presentations. This decrease in response is consistent with stimulus-specific adaptation.

### Omission Responses Encode Tone Probability

To further validate ORNs as negative prediction-error neurons, we investigated whether ORNs’ responses to omissions were modulated by the predictability of the omitted stimuli. If these neurons indeed encode negative prediction errors, we expect stronger omission responses in contexts where the tones are more predictable. This is because a more predictable stimulus generates a stronger predictive signal, which in turn leads to a more pronounced negative prediction error when the expected stimulus is omitted. Figure 4A illustrates the omission responses of three example ORNs across eight different conditions. These range from scenarios with 0% O_P_ tone (95% O_NP_ and 5% omission) to scenarios with 95% O_P_ tone (0% O_NP_ and 5% omission). A clear pattern emerges from these examples: omission responses intensified as the percentage of O_P_ tone in the sequence increased.

**Figure 4.**
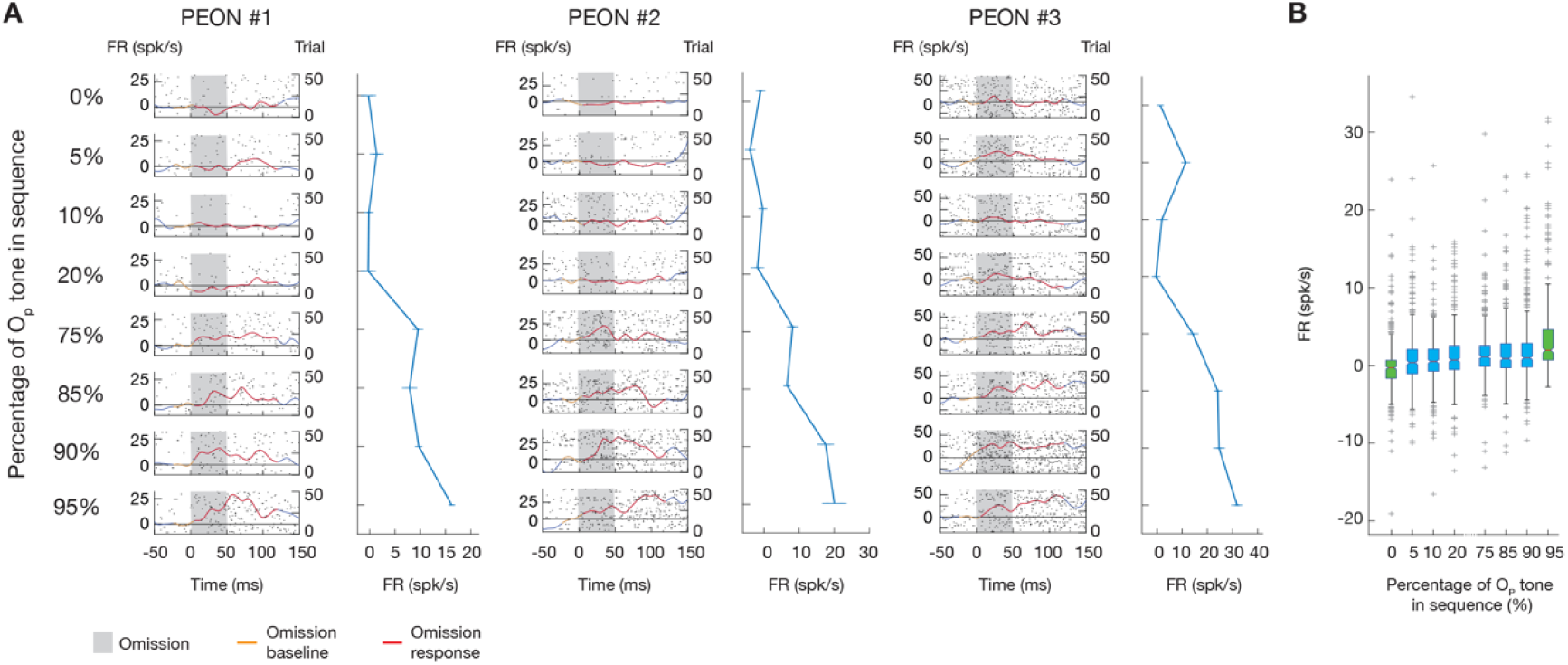
Probability encoding in ORNs. (**A**) Responses of three example PEONs to varying probabilities of the OP tone in the sequence. Raster plots and PSTH are shown for eight conditions with the percentage of the OP tone ranging from 0% to 95%. The x-axis represents time. The left y-axis represents the firing rate (FR), and the right y-axis represents the trial number. Gray shading indicates the omission period. The orange line represents the omission baseline, and the red line shows the omission response. The rightmost part of each panel shows the mean firing rate plotted against the percentage of the OP tone for each condition, with the blue line indicating the average firing rate across trials. (**B**) Box plot showing the firing rate of all PEONs across different probabilities of the OP tone. The x-axis represents the percentage of the OP tone in the sequence, and the y-axis represents the firing rate. Boxes represent the interquartile range (IQR), with the median indicated by a horizontal line. Notches indicate the 95% confidence interval for the median, allowing visual comparison of medians. Whiskers extend to 1.5 times the IQR, and outliers are shown as individual points, demonstrating that the firing rate of PEONs increases with higher probabilities of the OP tone. Green boxes highlight conditions with only one tone and omissions.

We then focused on ORNs that showed a significant correlation between their omission responses and the percentage of the O_P_ tone (as verified by a one-sided Spearman test, see details in Methods). Out of a total of 411 ORNs, we identified 143 that we have named “Probability Encoding Omission Neurons” (PEONs). Figure 4B presents a box plot that aggregates the omission response data across all PEONs. To quantify the relationship between omission response in PEONs and O_P_ tone probability, we employed a linear mixed-effects model with the omission response as the dependent variable, O_P_ tone probability as the fixed effect predictor, and random intercepts for each neuron, auditory tone, rat, and penetration. The model’s fixed effects revealed a significant effect of O_P_ tone probability on omission response (p < 1e-27, t (1142) = 11.238). Moreover, the adjusted R^2^ value of 0.4870 indicates that nearly half of the variance in omission responses among PEONs is explained by this model, further emphasizing the precision with which PEONs encode the likelihood of expected auditory events.

### Spatial Distribution of ORNs and PEONs in Auditory Cortex

To investigate the anatomical basis of predictive coding in the auditory system, we analyzed the laminar and areal distribution of ORNs and PEONs. We used current source density (CSD) analysis of LFP responses to determine cortical layer depths and categorized neurons into layers based on established criteria (see Methods). Figure 5 illustrates the distribution of these neurons across cortical depths. ORNs constituted a substantial fraction of neurons across all cortical layers, with the highest proportion in the granular layer. PEONs showed a different pattern, with higher representation in granular and supragranular layers compared to the infragranular layer. The laminar distribution of these neurons is summarized in Table 1.

**Figure 5.**
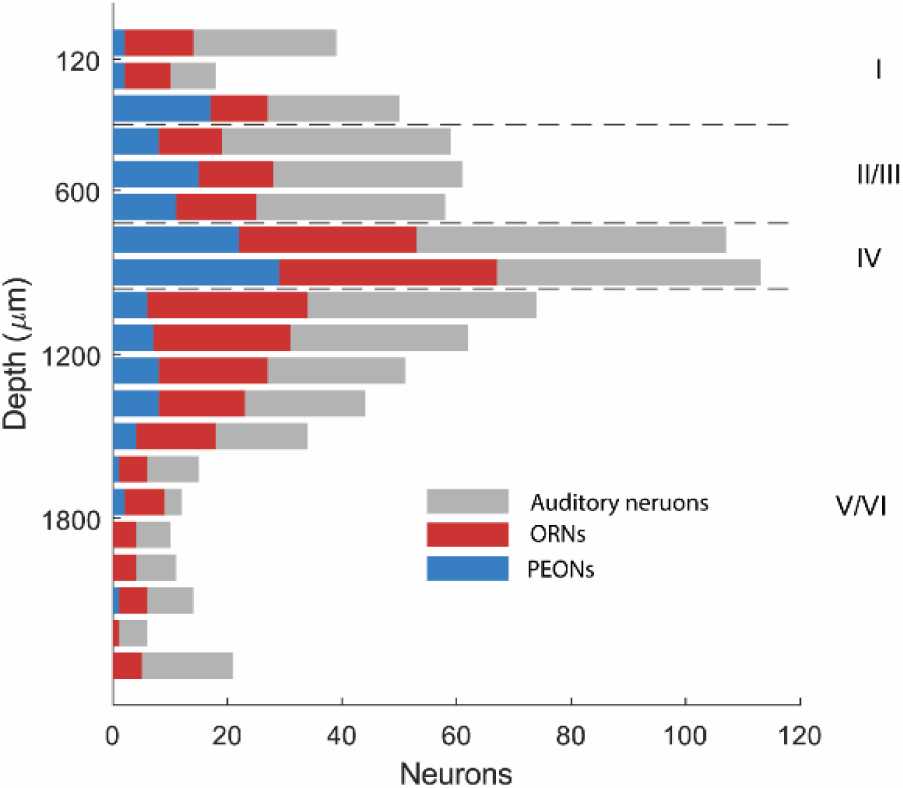
Laminar Distribution of Neurons in the Rat Auditory Cortex. Distribution of auditory neurons (gray), ORNs (blue), and PEONs (red) across cortical depths. The y-axis represents the cortical depth from the pial surface, with 0 μm corresponding to the cortical surface. The x-axis shows the normalized count of neurons. Horizontal dashed lines indicate the approximate boundaries between supragranular (0-600 μm), granular (600-900 μm), and infragranular (900-1400 μm) layers, based on current source density analysis.

**Table 1.** Distribution of Neurons Across Cortical Layers.

| Neuron Type | Supragranular (%) | Granular (%) | Infragranular (%) |
| --- | --- | --- | --- |
| All Auditory | 285 (100) | 232 (100) | 342 (100) |
| ORNs | 123 (43.16) | 127 (54.74) | 161 (47.08) |
| PEONs | 55 (19.30) | 52 (22.41) | 36 (10.53) |

To assess whether these distributions differed significantly from a random distribution, we performed a two-sided bootstrap analysis with 300,000 samples. For ORNs, the analysis revealed a trend towards a different distribution in the granular layer (p = 0.0851), although this did not reach statistical significance. For PEONs, we found a significant difference in distribution in the granular layer (p = 0.0140) and in the infragranular layer (p = 0.0004). Further examination of these significant results revealed a higher-than-expected presence of PEONs in the granular layer and a lower-than-expected presence in the infragranular layer. We further examined neuron distribution across different auditory fields, summarized in Table 2. Bootstrap analysis revealed no significant differences in the distribution of ORNs or PEONs across these areas (p > 0.05 for all comparisons), suggesting a consistent presence of these neurons throughout the auditory cortex.

**Table 2.** Distribution of Neurons Across Brain Areas.

| Neuron Type | A1 (%) | VAF (%) | AAF (%) |
| --- | --- | --- | --- |
| All Auditory | 534 (100) | 217 (100) | 108 (100) |
| ORNs | 257 (48.12) | 103 (47.47) | 51 (47.22) |
| PEONs | 94 (17.61) | 32 (14.75) | 17 (15.74) |

### PEONs Selectively Respond to Omissions While Broadly Reacting to Tones

As expected from the definition of PEONs, their omission response was highly selective. We observed that a substantial proportion of PEONs detected only the absence of one of the tones. 113 out of 139 PEONs (79%) showed a significant omission response during the sequence of 95% O_P_ tone and 5% omission but not during the sequence of 95% O_NP_ tone and 5% omission. This phenomenon is exemplified by the representative PEON shown in Figure 6A. Despite their selective omission response, PEONs generally responded to the presentation of both tones (O_P_ and O_NP_), as depicted in Figure 6B. This broad responsiveness is quantified by our finding that among the selective PEONs, 82 (or 73%) exhibited significant responses to both the O_P_ and O_NP_ tones across different experimental conditions (see details in Methods). Additionally, 13 PEONs responded exclusively to the O_P_ tone, 8 solely to the O_NP_ tone, and 10 did not significantly respond to either tone (see Figure 6C). This differential response pattern – selective to the omission of the O_P_ tone while broadly responsive to the presentation of both tones – highlights an intriguing asymmetry between top-down and bottom-up processing within the PEON population.

**Figure 6.**
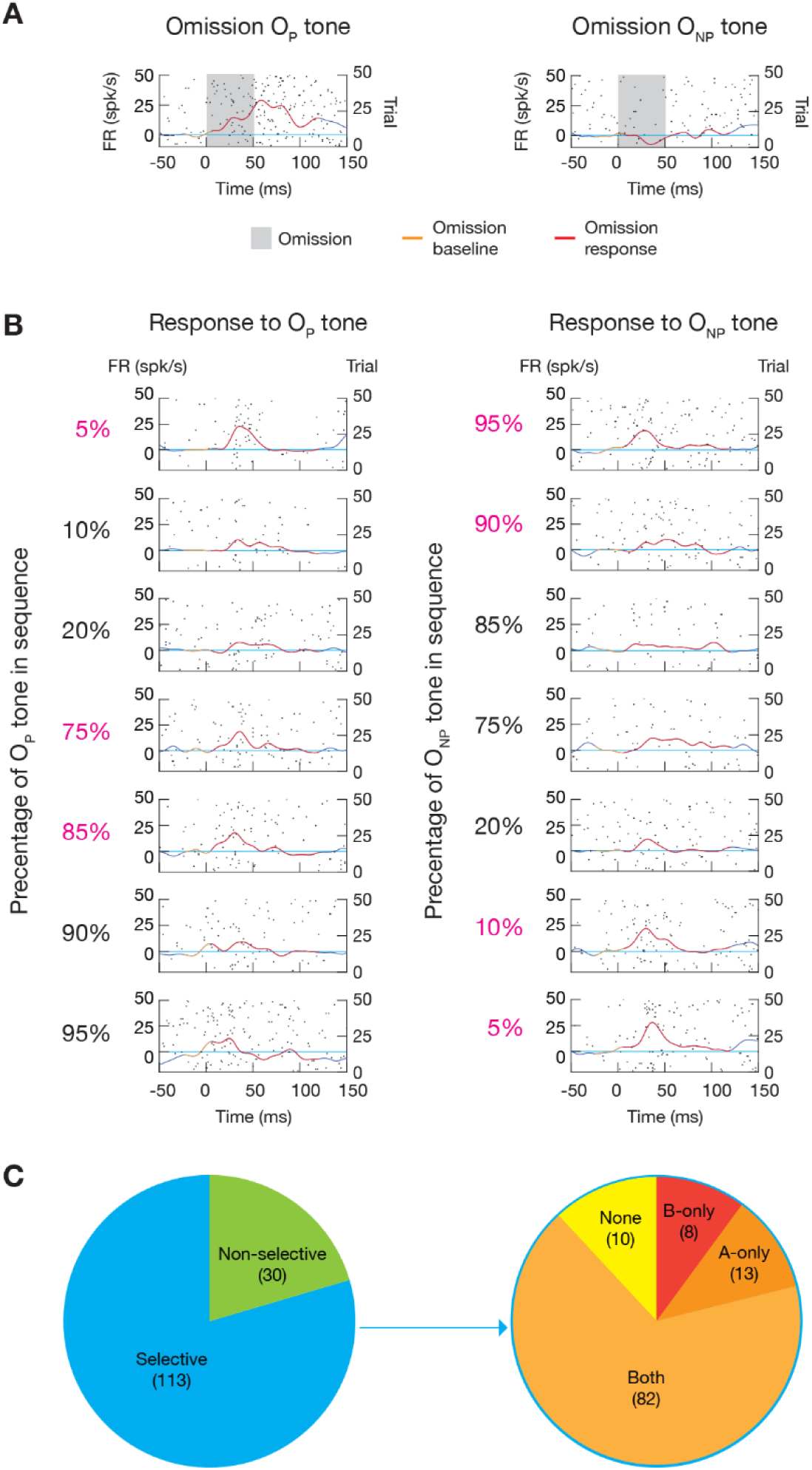
Selective omission and broad tone responses in PEONs. (**A**) Example PEON showing responses to the OP tone and ONP tone. The raster plot and PSTH show the firing rate over time. The left y-axis represents the firing rate, and the right y-axis represents the trial number. Gray shading indicates the omission period. The orange line represents the omission baseline, and the red line shows the omission response. (**B**) Responses to OP and ONP tones across varying probabilities of the tones in the sequence. The same representation is used as in panel A. The left column shows responses to the omission of the OP tone with different percentages of the OP tone in the sequence. The right column shows responses to the omission of the ONP tone with different percentages of the ONP tone in the sequence. The percentages in magenta indicate conditions with significant omission responses for the respective tones. (**C**) Pie charts summarizing the selectivity of PEONs. The left pie chart shows the distribution of selective and non-selective PEONs among the total identified PEONs. The right pie chart breaks down selective PEONs showing the number of neurons responding exclusively to the OP tone (A-only), exclusively to the ONP tone (B-only), both tones (Both), or neither tone (None).

### Neural Circuit Model for Asymmetric Predictive-Coding Processing

To further understand how PEONs, which we presume to be negative prediction-error neurons, can display this observed asymmetry in top-down and bottom-up processing, we developed a neural circuit model consisting of multiple streams within a “predictive-coding module” (Figure 7A). Each module includes six leaky integrate-and-fire (LIF) neurons divided into two streams: one stream calculates the positive prediction error and upregulates the prediction, while the other stream calculates the negative prediction error and downregulates the prediction. A sensory neuron (denoted as I) receives the sensory signal and propagates the information in a bottom-up manner. In contrast, a prediction neuron (P) receives the prediction signal and propagates the information in a top-down manner. A positive prediction-error neuron (PE+) receives excitatory synaptic input from the sensory neuron and is suppressed by the prediction neuron via an inhibitory interneuron (I+). Conversely, a negative prediction-error neuron (PE–) receives excitatory synaptic input from the prediction neuron and is suppressed by the sensory neuron via an inhibitory interneuron (I–). All synaptic connections are static synapses that include three variables: synaptic weight, time constant, and synaptic delay (see details in Methods). The activity of PE+ enhances the prediction signal through excitatory analog feedback, while the activity of PE– suppresses the prediction signal via inhibitory analog feedback.

**Figure 7.**
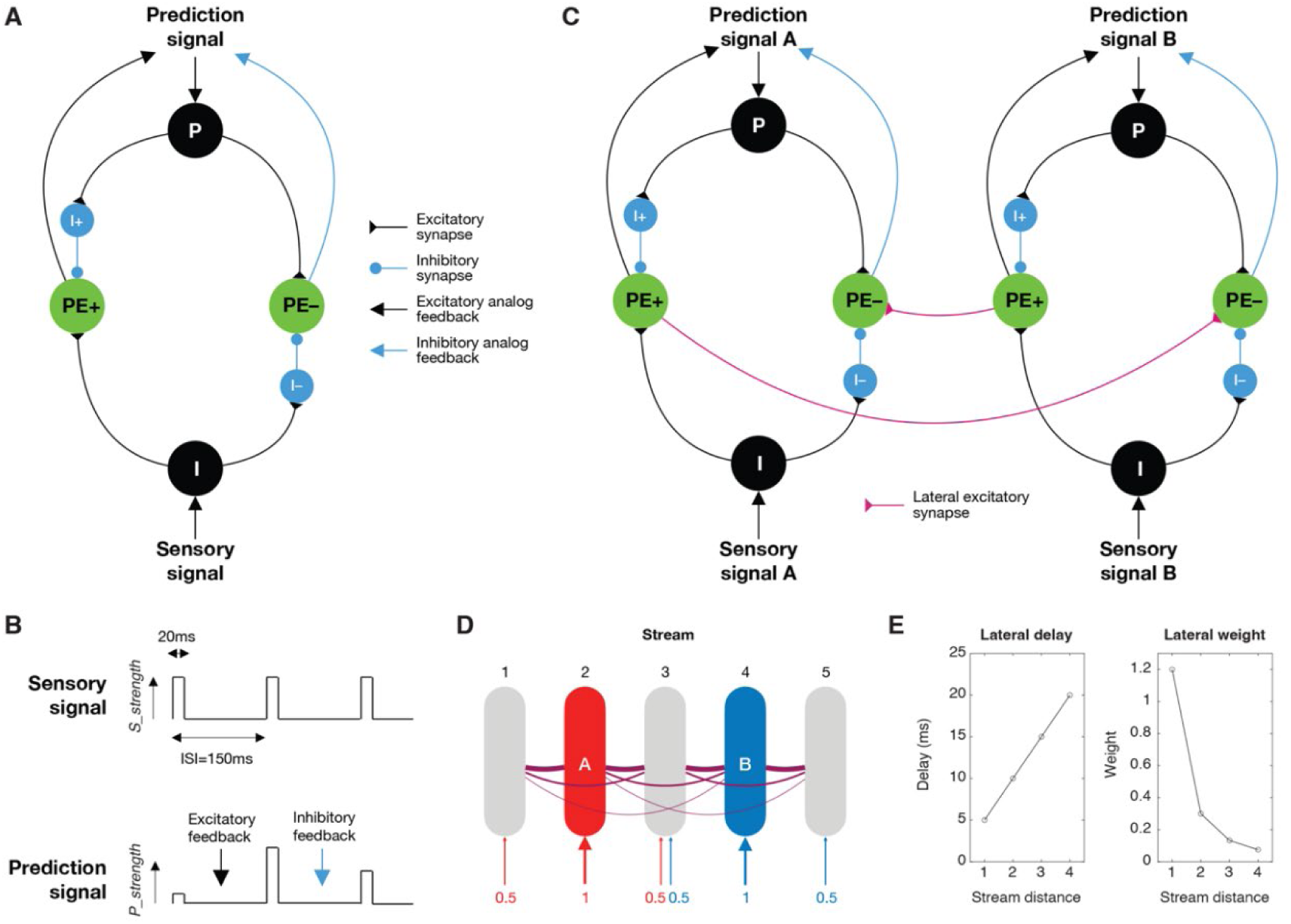
Neural circuit model for asymmetric predictive-coding processing. (**A**) Basic circuit module showing interconnections between neurons. P: prediction neuron, PE+: positive prediction error neuron, PE–: negative prediction error neuron, I+/I–: inhibitory interneurons, I: sensory input neuron. The connection types are also indicated. (**B**) Temporal profiles of sensory and prediction signals. (**C**) Lateral connections between two circuit modules processing different sensory streams. (**D**) Full model with 5 sensory streams showing tonotopic organization. Streams 2 and 4 receive full strength input for tones A and B respectively. The numbers below illustrate the strength of sensory signal (*S_strength*) in each stream. (**E**) Distance-dependent properties of lateral connections, showing delay and weight as a function of the distance between streams (see more details in Methods).

In our simulation, the sensory signal was designed to replicate the experiment, featuring an analog pulse stimulus with an intensity of *S_strength* (ranging between 0 and 1) and a duration of 20ms, delivered with an inter-stimulus interval (ISI) of 150ms (see Figure 7B, top). Conversely, the prediction signal was modeled as a similar analog pulse as the sensory signal, but varied in intensity (*P_strength*), which could be enhanced by excitatory feedback from PE+ or diminished by inhibitory feedback from PE– (Figure 7B, bottom). It is important to recognize that our model assumes the temporal profile of the prediction signal is learned through a separate mechanism, while the predictive-coding module adjusts only the signal’s magnitude based on prediction error.

Predictive coding modules from different streams, each processing a specific frequency according to a tonotopic organization, are interconnected through lateral excitatory synaptic connections. These connections run from PE+ of one module to PE– of another (as illustrated between streams A and B in Figure 7C). Intuitively, this setup enables PEONs, represented as PE–, in one stream to respond to stimuli in another stream, endowing them with a broader bottom-up receptive field. Moreover, this configuration allows an unexpected stimulus in one stream to inhibit the predictive signal in other streams. For example, as shown in Figure 7C, if the sensory signal in Stream B is unexpected, PE+ in Stream B will activate, causing PE– in Stream A to also activate through the lateral connection. This activation will then suppress the predictive signal in Stream A. Thus, when an unexpected Tone B is detected, indicating that Tone A (or other tones) is unlikely, the prediction for Tone A (or other tones) is consequently reduced. Conversely, since the absence of an expected Tone B does not necessarily suggest the presence of Tone A (or other tones), our model does not include lateral connections running from PE– to PE+.

Our comprehensive model includes five streams: Stream 2 is tuned to detect the O_P_ tone (Tone A), while Stream 4 detects the O_NP_ tone (Tone B) (see Figure 7D). The model incorporates a simple receptive field structure for sensory signals. Specifically, Stream 2 receives Tone A at full strength (*S_strength*=1), whereas the adjacent Streams 1 and 3 receive Tone A at half strength (*S_strength*=0.5). Similarly, Stream 4 receives Tone B at full strength, and the adjacent Streams 3 and 5 receive Tone B at half strength. Additionally, the model features lateral excitatory connections between streams that are distance-specific (Figure 7E). The synaptic delay between any two streams is proportional to their spatial separation, and the synaptic weight decreases with the inverse square of their distance (see more details in Methods).

### Asymmetric Activities in Modelled PEON

We conducted multiple simulations using the model, each featuring unique probability combinations of tones A and B, along with a consistent 5% omission rate, mirroring the conditions of our rodent experiments. Each simulation included 500 stimuli—either tone A, B, or an omission (O)—across a total of 75 seconds. Importantly, every simulation began with no initial predictions across all streams (*P_strength*=0).

Here, we demonstrate the model’s responses using the example where the sensory inputs consist of 90% A, 5% B, and 5% O, randomly ordered as shown in Figure 8A. The spiking activities of Neurons I, P, PE+, and PE− across the five streams are illustrated in Figure 8B. Notably, Neuron P began firing after a few seconds, particularly in Streams 1-3, which are responsive to Tone A. This activity indicated that the prediction for Tone A was beginning to be established. Consequently, the positive prediction error, reflected by PE+’s spiking activity, began to diminish. Additionally, the spiking activity of PE−, primarily associated with the omission of Tone A, was also observed in Streams 4 and 5. This was largely due to lateral connections from PE+ in Streams 1-3. The prediction signals across the five streams, detailed in Figure 8C, show that *P_strength* increased and plateaued in Streams 1-3, caused by PE+’s firing, with Stream 2 displaying the highest value.

**Figure 8.**
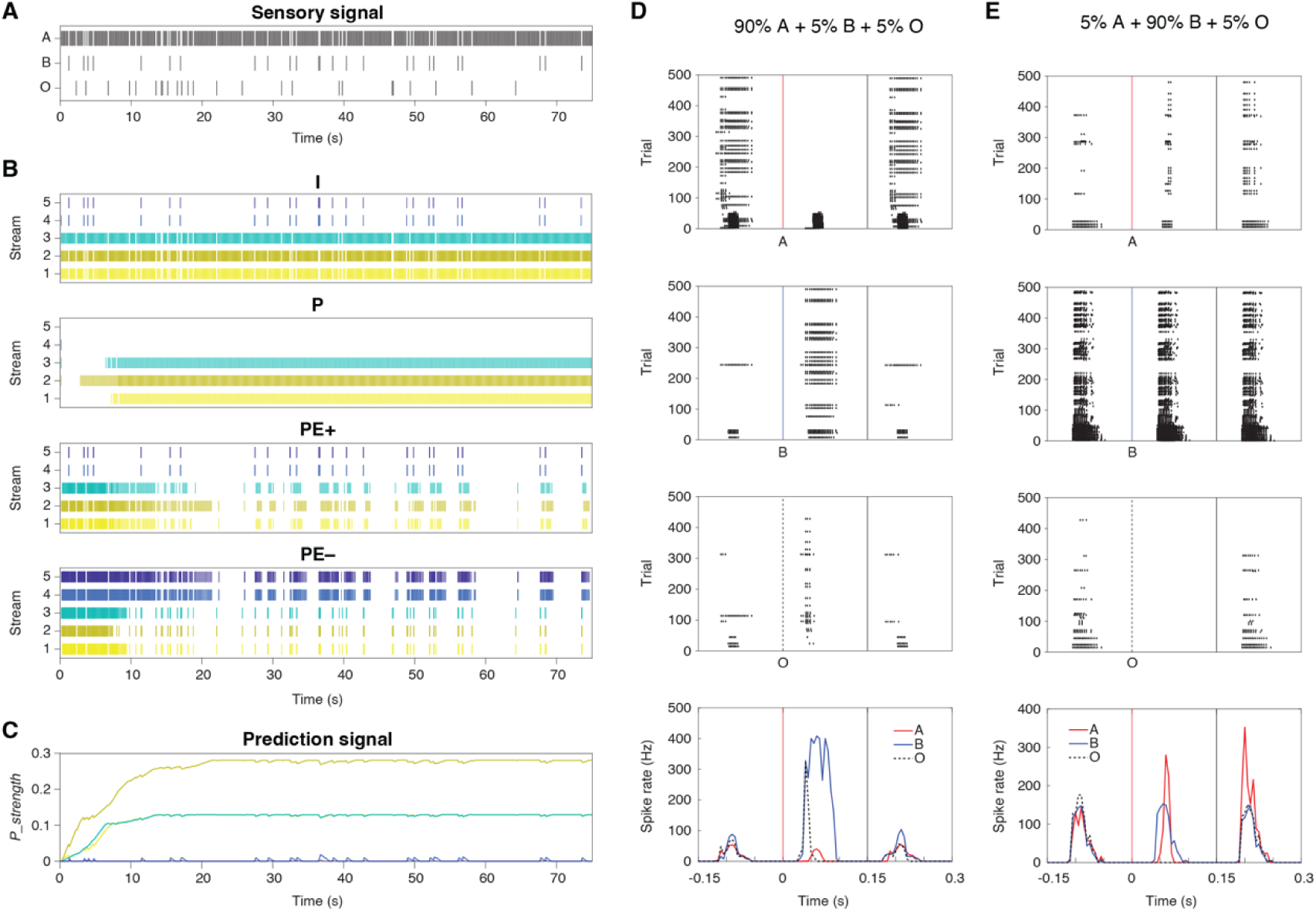
Asymmetric activities in modelled PEON. (**A**) Sensory input sequences over 75 seconds, showing occurrences of tones A (90%), B (5%), and omissions (5%). (**B**) Spiking activities of neurons I, P, PE+, and PE– across five streams. Colors represent different streams. (**C**) Prediction signal strength (*P_strength*) over time for different streams. The same color scheme is used as in panel B. (**D**) Raster plots (top 3 panels) and PSTHs (bottom panel) for PEON in Stream 2 under: 90% A, 5% B, 5% O. Responses to tones A (red vertical line), B (blue vertiacal line), and omissions (O, black dotted vertical line) are shown separately. PSTHs display firing rate over time, with colored lines indicating responses to different stimuli. (**E**) Raster plots and PSTHs for PEON in Stream 2 under: 5% A, 90% B, 5% O.

To explore asymmetric predictive-coding processing, we focused on the PEON in Stream 2, the target neuron for modeling our experimental results. Our goal was to reproduce the experimentally observed asymmetry: selective responses to omissions of the preferred tone, but broad responses to both tones. The raster plots of this PEON, presented in Figure 8D for a simulation containing 90% A, 5% B, and 5% O, characterize responses to different stimuli: Tones A, B, and omissions O. The post-stimulus time histogram (PSTH) reveals that while the PEON consistently responded to omissions, it did not respond to Tone A (in later trials), which was anticipated. Importantly, the PEON did respond to Tone B, a response enabled by lateral connections. In Figure 8E, we present raster plots from a different simulation with 5% A, 90% B, and 5% O. Similar to the previous simulation, the PEON in Stream 2 responded to Tone B via lateral connections. It also reacted to Tone A, facilitated by the lateral connections from nearby Streams 1 and 3, where PE+ was activated by the deviant Tone A. Notably, there was no response to omissions, as the prediction in Stream A was not effectively established due to the minimal occurrence of only 5% A. In conclusion, the PEON in Stream 2 demonstrated the asymmetry of receptive fields in bottom-up sensory and top-down predictive processing, reflecting the findings in the rodent experiment.

### Alternative Models and Evaluation

To assess the validity of the proposed model with lateral connections (the model labeled “Lateral PE+ to PE–“), we evaluated two alternative models and compared their predictions against the experimental data. The alternatives included: a model with lateral connections from Neuron I to PE– across different streams (termed “Lateral I to PE–“, depicted in Figure 9A), and a model with no lateral connections at all (referred to as “No lateral”). For each model, we measured the responses of the PEON in Stream 2 to Tones A, B, and omission O under different probabilities of Tone A (Prob(A)): from 0% (95% B and 5%O) to 95% (0% B and 5%O). The results are shown in Figure 9B, together with the correlation between the firing rate and Prob(A).

**Figure 9.**
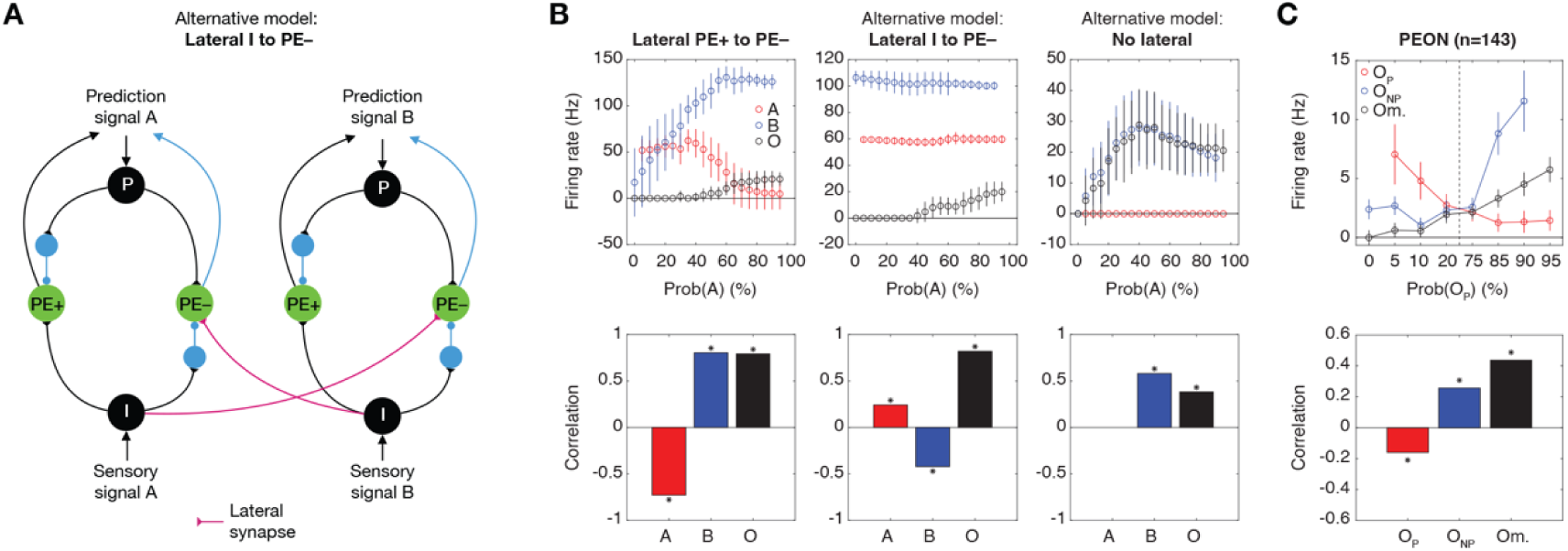
Alternative models and evaluation. (**A**) Schematic of the alternative model with lateral connections from input neurons (I) to negative prediction error neurons (PE–) across streams A and B. (**B**) Evaluation of alternative models: Lateral PE+ to PE-, Lateral I to PE-, and No Lateral. Top panels show the firing rates of PEON in Stream 2 in response to Tone A (red), Tone B (blue), and omissions (O, gray) across varying probabilities of Tone A (Prob(A)). Bottom panels display correlations between firing rates and probabilities for Tones A, B, and omissions for each model. (**C**) Experimental validation with PEONs (n=143). Top panel shows firing rates of PEONs in response to OP tones (blue), ONP tones (red), and omissions (Om., gray) across varying probabilities of OP tones (Prob (OP)). Bottom panel displays correlations between responses to OP tones, ONP tones, and omission and OP probabilities .

In our proposed model, the PEON’s response to omission significantly correlated with Prob(A) (r= 0.7912, p= 2.1e-108, Spearman rank coefficient, n= 500 trials; see Methods for further details), indicating that the PEON did encode the probability of the stimulus within its stream. For Tone A, the PEON exhibited reduced responses as Prob(A) increased, demonstrating a significant negative correlation between firing rate and Prob(A) (r= –0.7290, p< 1e-250, n= 4750 trials). This occurred because PE+ in nearby Streams 1 and 3 detected less surprise at higher Prob(A) values, leading to less activation of the PEON in Stream 2 through lateral connections (as also noted in Figure 7E). Conversely, for Tone B, the PEON’s response positively correlated with Prob(A) (r= 0.8051, p< 1e-250, n= 4750 trials). At higher Prob(A) values, Tone B emerged as the deviant tone. This led to an enhanced detection of surprise by PE+ near Stream 4 for Tone B, which subsequently activated the PEON in Stream 2 more strongly through lateral connections.

In the alternative “Lateral I to PE–” model, the PEON’s response to omission was also correlated with Prob(A) (r= 0.8206, p= 4.3e-123, n= 500 trials). For Tone A, unlike the proposed model, the PEON in Stream 2 did not exhibit decreased responses at high Prob(A) values. This occurred because the PEON directly received sensory input from Neuron I, which remained unchanged across different Prob(A) settings. Despite this, a positive correlation between the firing rate and Prob(A) was observed (r= 0.8206, p= 4.3e-123, n= 4750 trials). This was due to slightly increased responses from Neuron I in nearby Streams 1 and 3, which were caused by an elevated membrane potential resulting from frequent stimulation by Tone A. Conversely, for Tone B, the opposite effect occurred: Neuron I near Stream 4 showed an increase in membrane potential due to frequent stimulation of Tone B at low Prob(A) levels. Therefore, despite only minor changes in firing rate across different Prob(A) values, a negative correlation was observed (r= –0.4217, p= 3.5e-204, n= 4750 trials).

In the alternative “No lateral” model, the PEON’s response to omission correlated with Prob(A) (r= 0.3814, p= 9.2e-19, n= 500 trials), as this encoding occurs within the stream without the need for lateral connections. For Tone A, the PEON displayed no responses, which was anticipated since it could only react to an omission in the absence of lateral connections. For Tone B, the PEON exhibited similar responses as to omissions, since in Stream 2, any stimulation differing from Tone A was treated as an omission. Consequently, a similar positive correlation was observed (r= 0.5814, p< 1e-250, n= 4750 trials).

To validate these models, each demonstrating unique encoding characteristics, we compared them to data from our rodent experiment. In our analysis, Tones A and B corresponded to the O_P_ and O_NP_ Tones, respectively. The correlation analysis indicated a positive correlation for omission responses (r= 0.4379, p= 8.7e-55, Spearman rank coefficient, n= 1144 trials; details in Methods), a negative correlation for responses to O_P_ tone (r= –0.1593, p= 4.0e-7, n= 910 trials), and a positive correlation for responses to O_NP_ tone (r= 0.2573, p= 1.3e-16, n= 910 trials) (Figure 9C). These results support the proposed model with its lateral connections facilitating positive prediction errors, distinguishing it from the alternative models.

The models discussed so far have not taken into account that the strength of sensory signals could diminish due to frequent exposure to a specific stimulus. To address this, we analyzed responses from neurons in the thalamus (n= 130 neurons) across various probabilities of the O_P_ Tone (see Supplementary Figure S1A and Methods for details). Accordingly, we modeled the sensory signal strength, *S_strength*, as a function of Prob(A) (Figure S1B) and conducted simulations. Our results reaffirmed that only the proposed model successfully replicated the probability encoding characteristics observed in the rodent data (Figure S1C).

### Computational Benefits of Lateral Connections

Our next step was to explore the potential computational benefits of the proposed lateral connections, particularly in terms of enhancing prediction encoding. How do these connections influence prediction signals? Without lateral connections, not only would Stream 2 predict Tone A, but also the neighboring streams (e.g., Stream 1), due to their broad sensory receptive fields receiving half-strength sensory signals. However, with lateral connections, these predictions are reduced not only by the omission of Tone A but also by the presence of Tone B. Particularly, we theorized that this additional reduction would suppress the already low prediction signals in the neighboring streams. This leads to a scenario where the prediction signal is predominantly strong in Stream 2, thus resulting in a more precise encoding of Prob(A) across streams.

To test this, we first assessed the steady value of *P_strength* after the prediction signal stabilized across various stimulation sequences (as in Figure 8C). These sequences varied Prob(A) from 0% to 95%, with increments of 1%. In Figure 10A, we plotted the time courses of *P_strength* for each 75-second simulation under different Prob(A) settings, focusing on Stream 2, where our target PEON was situated, and the adjacent Stream 1. For comparison, the time courses for Streams 1 and 2 in the model without lateral connections (“No lateral”) are shown in Figure 10B. The steady value of *P_strength* was determined by averaging values over the last 50 seconds (from 25 to 75 seconds). It was plotted as a function of Prob(A) for both models, with and without lateral connections, across Streams 1 and 2 (as shown in Figure 10C). In both streams, *P_strength* increased as Prob(A) rose. However, the increase was less pronounced in the model with lateral connections, especially at lower Prob(A) values. We further analyzed the contrast in *P_strength* between Streams 1 and 2 by calculating the decibel value (logarithmic ratio of *P_strength* in Stream 2 to that in Stream 1). This contrast was significantly greater in the model with lateral connections compared to the model without, particularly when Prob(A) was less than 0.5 (Figure 10D). This suggested that the lateral connections enhance the contrast of prediction signals across streams.

**Figure 10.**
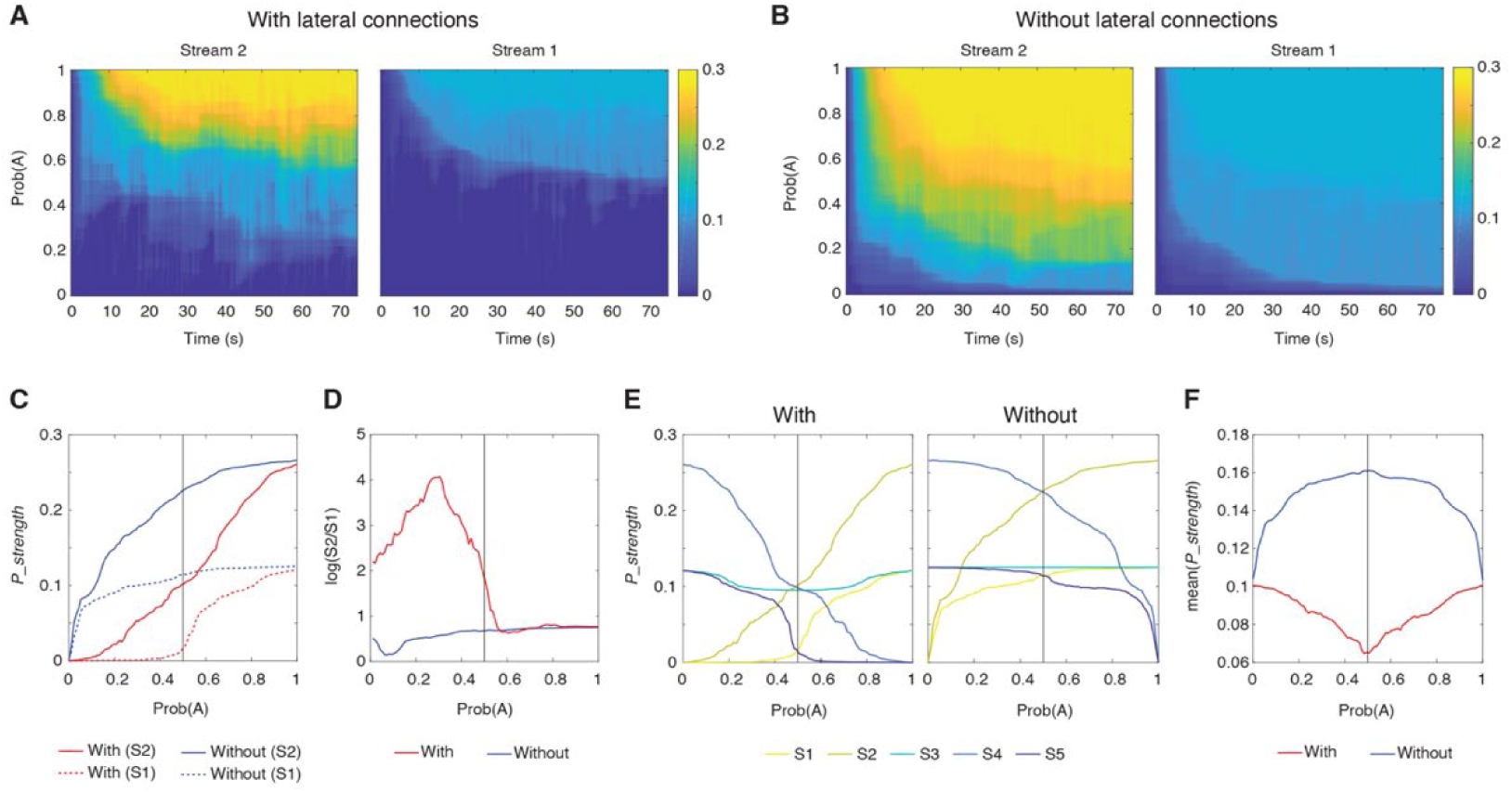
Computational benefits of lateral connections. (**A**) Time courses of prediction strength (*P_strength*) in Streams 2 and 1 with the lateral connections for varying probabilities of Tone A. Colors indicate the magnitude of *P_strength*. (**B**) Time courses of prediction strength in Streams 2 and 1 without the lateral connection. The same representation is used as in the panel A. (**C**) Steady-state *P_strength* in Stream 2 (solid lines) and Stream 1 (dotted lines) as a function of Prob(A) with (red) and without (blue) lateral connections. (**D**) Contrast in *P_strength* between Streams 1 and 2 (log scale) with and without lateral connections, plotted against Prob(A). (**E**) Steady-state *P_strength* across all streams (S1 to S5) as a function of Prob(A), with and without lateral connections. (**F**) Mean *P_strength* across all streams for different Prob(A) settings, comparing the network with (blue) and without (red) lateral connections.

We also hypothesized that the additional suppression of prediction signals facilitated by lateral connections could enhance prediction encoding efficiency. This is supported by the observation that *P_strength* was generally lower with lateral connections (as shown in Figure 10C). To investigate this further, we measured *P_strength* across Prob(A) for all streams in both models (Figure 10E), and then calculated the average *P_strength* across streams for each Prob(A) setting (Figure 9F). The findings suggest that with lateral connections, the network requires less “energy” or smaller prediction signals to effectively encode sequences, particularly in less predictable scenarios where the occurrences of Tones A and B are equal, such as Prob(A) ≈ 0.5. This demonstrates a more efficient encoding strategy. Conversely, without lateral connections, the network expends more energy to manage more unpredictable sequences.

### Lateral Prediction Suppression and the Free Energy Principle

The free energy principle provides a powerful theoretical framework for understanding how the brain processes information and generates predictions about the world [31]. In this section, we demonstrate how our experimental findings on lateral prediction suppression could naturally emerge from the free energy principle, offering deeper insight into the computational mechanisms underlying auditory processing. The complete mathematical derivations and detailed equations are provided in Supplemental Information, allowing us to focus here on the key concepts and implications.

We consider the simplest form of the free energy principle, where the parameters of the model, including prior expectations of hidden variables and covariance matrices, are assumed to be adapted to the natural environment and stable over the time scale of the experiments. We also consider only one hierarchy in the predictive coding network, mirroring our generative model’s structure, and only two tone frequencies for the sensory input, as in our electrophysiological experiments. In such a case, we derive a prediction update rule comprising three terms (see Supplemental Information, Eq. 11). The first term represents the update by the positive and negative prediction errors in the same stream. The second term represents the lateral interaction of positive and negative prediction errors across steams. The third term represents the temporal decay of the prediction. Thus, the lateral interaction of positive and negative errors across different streams is a natural outcome derived directly from the free energy principle.

## Discussion

This study aimed to understand how negative prediction-error neurons in the rat auditory cortex respond. We combined electrophysiological recordings with computational modeling to demonstrate an asymmetry in prediction-error signaling, which we attribute to “lateral prediction suppression” where positive and negative prediction errors interact through lateral connections to enhance the precision and efficiency of prediction encoding.

### Identification of Omission-Responsive Neurons

Omission paradigms have long been employed to identify neurons responsible for encoding negative prediction errors. In human studies using EEG and MEG, mismatch negativity (MMN) responses to unexpected omissions in tone sequences have been reported, which are interpreted as evidence for predictive coding mechanisms [17,20,22–24,32]. However, until recently, there was limited evidence for omission responses at the single-neuron level in animal models. Recently, Awwad et al. [25] demonstrated robust omission responses in unanesthetized rats using complex stimuli involving the omission of expected gaps in auditory stimulation. Despite providing compelling evidence for prediction-related responses, the complexity of their stimuli poses challenges for direct comparisons with studies employing simpler omission paradigms.

Lao-Rodríguez et al. [26] reported significant omission responses in 35.6% of neurons in awake animals and 19.5% in anesthetized rats and mice. In contrast, our study identified a higher percentage of omission-responsive neurons (48% in anesthetized rats). This discrepancy may be attributed to several factors: our use of sequences with varying probabilities likely increased the detection of neurons responding selectively to tone omissions; the higher sensitivity and fidelity of Neuropixels electrodes compared to traditional methods; and the ability to record from many neurons simultaneously, allowing us to detect omission-responsive neurons with weaker or more selective responses to standard stimuli.

### Mechanistic Insights and Comparison to Existing Models

Our model builds upon the framework proposed by Keller and Mrsic-Flogel [6], which describes predictive coding as a canonical cortical computation. Their model emphasizes hierarchical processing with distinct neuron populations computing positive and negative prediction errors. Our study provides empirical evidence for these mechanisms in the auditory cortex, extending previous findings on negative prediction error neurons in visual cortex [7,9] to the auditory domain. Another key advance in our model is the incorporation of lateral connections between prediction-error neurons across adjacent frequency streams. This mechanism enables complex interactions, refining prediction signals and addressing the asymmetry between selective omission responses and broad tone responses. By capturing the response properties of PEONs and their ability to encode stimulus probability, our study extends the predictive coding framework across sensory modalities.

### Lateral Connections in Predictive Coding Networks

Our circuit model shows that unexpected stimuli in one stream inhibit predictive signals in adjacent streams, reducing prediction noise and enabling rapid updates to internal models. While similar to lateral inhibition in the visual system [33], our mechanism involves excitatory connections that refine predictions. Biological evidence strongly supports the existence of excitatory lateral connections between adjacent frequency channels in the primary auditory cortex (A1). These connections play a crucial role in spectral integration and the processing of complex auditory stimuli [34–36]. Anatomical studies have revealed that pyramidal neurons in A1 exhibit long-range horizontal collaterals that form connections across different frequency representations, often aligning with isofrequency bands [37,38]. Physiological investigations have demonstrated that these intracortical pathways contribute significantly to the breadth of subthreshold frequency receptive fields, which can span up to five octaves [39,40]. The strength and probability of these connections are distance-dependent, with stronger and more probable connections between neurons tuned to close frequencies, weakening as the frequency distance increases [41,42]. This organized network of lateral connections facilitates the integration of spectral information across different frequency channels, supporting complex auditory processing and cortical plasticity in A1.

Similar patterns of lateral connectivity have been observed in other sensory modalities, such as orientation-specific connections in visual cortex [43] and whisker-related connections in somatosensory barrel cortex [44], suggesting a common organizational principle across different sensory areas. The development of these lateral connections might involve both innate and learned components. While the basic framework of these connections may be genetically predetermined, their refinement occurs during early postnatal development through experience-dependent plasticity [45]. This developmental process allows for the fine-tuning of these connections based on sensory experience, optimizing the auditory cortex for processing the specific acoustic environment encountered during critical periods of development.

In our model, the one-way excitation from PE+ to PE-neurons allows for rapid recalibration of expectations across streams. This asymmetry enables efficient updating of predictions based on positive surprises while avoiding false alarms from mere absences. Integration across frequency channels enables complex interactions, with sequences of tones providing predictive information about subsequent tones. Naturalistic sounds like tone clouds can activate multiple streams simultaneously, leveraging the network’s capacity for sophisticated auditory processing [46].

### Layer and Area Specificity of PEONs

Our finding that PEONs are more prevalent in granular and supragranular layers aligns with current views on the laminar organization of predictive coding circuits [4]. These layers are thought to be primary sites for prediction error computation and integration of top-down predictions with bottom-up sensory input. The relative scarcity of PEONs in infragranular layers suggests these neurons may be less involved in generating feedback predictions to earlier processing stages. The even distribution of PEONs across different auditory fields (A1, VAF, AAF) suggests that predictive processing is a fundamental computation throughout the auditory cortex, consistent with the idea of predictive coding as a canonical cortical algorithm [6]. However, this raises questions about potential functional specializations not captured by our current paradigm.

### Establishment of Prediction Signals

A notable limitation of our study is the absence of explicit “prediction neurons” in both our experimental findings and computational model. While we identified neurons responding to prediction errors, we did not find neurons directly representing predictions about upcoming stimuli. Predictive signals might be generated in higher-order auditory areas, non-auditory regions providing top-down input, or encoded in a distributed manner across neuronal populations. Further computational modeling of distributed prediction encoding could help guide future experiments by suggesting specific patterns or population-level dynamics to look for in neural data.

Future research should address the limitations of our study, such as the potential effects of anesthesia on neural responses, by comparing findings with those from awake and behaving animals. Additionally, employing more complex stimuli and varying spectral distances between tones could further probe the limits of the proposed lateral prediction suppression mechanism.

## Methods

### Animals

All experimental procedures were conducted in strict accordance with the ethical guidelines established by the Japanese Physiological Society for the care and use of animals in physiological studies. The experimental protocol was approved by the committee on the ethics of animal experiments at the Graduate School of Information Science and Technology, the University of Tokyo (JA21-9). The study involved ten male Wistar rats, aged 11-12 weeks, and weighing between 250 and 350 grams. The animals were housed in a carefully controlled environment with a 12-hour light/dark cycle, ensuring optimal conditions for their well-being. Food and water were provided ad libitum throughout the study. To minimize animal suffering, all surgical procedures were conducted under appropriate anesthesia. Upon completion of the experiments, the animals were humanely euthanized using an overdose of pentobarbital sodium (160 mg/kg, administered intraperitoneally), following established guidelines for the humane termination of animal studies.

### Surgical procedures

The rats were anesthetized using urethane, with an initial dose of 1.5 g/kg administered intraperitoneally and supplementary doses of 0.5 g/kg provided as needed to ensure stable and deep anesthesia. This depth of anesthesia was confirmed by the absence of corneal and pedal reflexes. The animals were secured using a custom-made head-holding apparatus. Atropine sulfate (0.1 mg/kg) was administered subcutaneously at the onset of the procedure to reduce bronchial secretion viscosity. Local anesthetic xylocaine (0.3–0.5 ml) was applied at the incision site before making the skin incision. A needle electrode was inserted subcutaneously into the right forepaw to serve as ground. A small craniotomy was performed near the bregma landmark to place a reference electrode, ensuring electrical contact with the dura. A larger craniotomy was performed over the right auditory cortex (4.0 mm posterior and 6.0 mm lateral to bregma), and the right temporal muscle, cranium, and dura covering the auditory cortex were surgically removed. The exposed cortical surface was kept moist with saline to prevent drying. Cisternal cerebrospinal fluid drainage was conducted to minimize cerebral edema. Throughout the experiment, an adequate and stable level of anesthesia was ensured by monitoring the animals’ respiratory rate, heart rate, and hind-paw withdrawal reflexes.

### Electrophysiological recordings

Neuronal activity was recorded using both a custom-designed surface microelectrode array[47] and a Neuropixels 1.0 electrode array (IMEC, Belgium)[48].

#### Mapping auditory cortex using the surface array

To record neuronal activity epipially over the auditory cortex, a surface array (NeuroNexus, Ann Arbor, MI, USA) comprising sixty-four platinum electrodes was employed. Each electrode had a diameter of 100 μm and a center-to-center distance of 500 μm, covering an area of 4.5 × 3.0 mm. The array featured 0.3-mm diameter holes between the electrodes to allow the insertion of a Neuropixels array. Auditory evoked potentials (AEPs) were recorded using amplifiers with a gain of 1000, a digital filter bandpass of 0.3–500 Hz, and a sampling frequency of 1 kHz, employing the Cerebus Data Acquisition System (Blackrock Microsystems LLC, Salt Lake City, UT, USA).

We mapped the click-evoked responses to pinpoint the location of the auditory cortex (AC). A click was defined as a monophasic positive wave lasting 20 ms. Sixty clicks were administered at a rate of 1 Hz, and local field potentials (LFPs) were recorded. The peak amplitude of the middle-latency auditory-evoked potential (P1) within the first 50 ms following the stimulus onset was measured from the grand averaged LFP. To identify three subfields within the AC—the primary auditory cortex (A1), the anterior auditory field (AAF), and the ventral auditory field (VAF)—we analyzed the spatial distribution of the P1 response.

#### Neuropixels recordings

For recording neuronal activity across different cortical layers, we used the Neuropixels 1.0 electrode array[48]. The array features 384 active recording sites (out of 960 sites total) located at the bottom ∼4 mm of a ∼10 mm shank (70 µm wide, 24 µm thick, 20 µm electrode spacing), with the reference and ground, shorted together. The probe’s ground and reference pads were soldered to a pin, which was then connected to a skull screw on the animal. Recordings were made in internal reference mode.

Data acquisition for the Neuropixels arrays was performed using Neuropixels version 1.0 acquisition hardware, specifically the National Instruments PXIe-1071 chassis and PXI-6133 I/O module for recording analog and digital inputs (Neuropixels version 1.0, IMEC, Belgium). The signal was collected from the most distal 384 recording sites (bank 0) at a 30 kHz sampling rate. Raw voltage traces from the Neuropixels arrays were filtered, amplified, multiplexed, and digitized on-probe. Data acquisition and initial processing were managed using Open Ephys software (www.open-ephys.org). Spike sorting was conducted using Kilosort 3 [49], and further manual curation was performed with Phy software (https://phy.readthedocs.io/en/latest/).

#### Characterization of the frequency response area

To assess the frequency-tuning properties of all recorded neurons, we measured the frequency response area (FRA) at each recording site. The auditory stimuli consisted of pure tone bursts with a 5-ms rise/fall time and a 20-ms steady-state duration. The tones covered a wide range of frequencies, spanning 18 distinct values from 1.6 to 64 kHz, and were presented at seven different sound pressure levels, ranging from 20 to 80 dB SPL in 10-dB steps. In total, 126 unique tone bursts were used to characterize the FRA. The stimuli were delivered in a pseudorandom sequence with an inter-tone interval of 600 ms, and each tone was repeated 20 times. FRA maps were generated for each neuron based on their responses to the presented tones. After examining the FRA maps, we selected two tones that were one octave apart and elicited robust responses from most recorded neurons. These two tones were then used as the stimuli for the subsequent experiments investigating the neural responses to tone sequences and omissions using an intensity well above the threshold of a typical neuron.

## Experimental Design

To investigate the neural responses to tone sequences and omissions, we designed eight experimental conditions, each employing an oddball paradigm with two distinct tones and a fixed 5% probability of tone omissions. The stimuli were the two sinusoidal tones chosen before with a duration of 50 ms, including 5-m rise and fall ramps. These tones were arranged in sequences of 1000 stimuli, with a stimulus onset asynchrony (SOA) of 150 ms. In the first condition, one tone (Tone A) comprised 95% of the sequence, while the other tone (Tone B) was absent. Across the subsequent conditions, the proportion of Tone B gradually increased in 5% increments, while the proportion of Tone A decreased accordingly. This resulted in the following sequences:

Sequence 1: 95% Tone A, 0% Tone B, 5% omissions

Sequence 2: 90% Tone A, 5% Tone B, 5% omissions

Sequence 3: 85% Tone A, 10% Tone B, 5% omissions

Sequence 4: 75% Tone A, 20% Tone B, 5% omissions

Sequence 5: 50% Tone A, 45% Tone B, 5% omissions

Sequence 6: 20% Tone A, 75% Tone B, 5% omissions

Sequence 7: 10% Tone A, 85% Tone B, 5% omissions

Sequence 8: 0% Tone A, 95% Tone B, 5% omissions

This design allowed us to systematically manipulate the relative probabilities of the two tones while maintaining a consistent rate of omissions. By progressively shifting the balance between the tones across conditions, we could examine how the neural responses to tone omissions were influenced by the statistical context provided by the sequence.

## Data analysis

### Auditory neuron classification

Using the responses to the tone sequences and omissions, we defined auditory neurons based on their activity patterns. To classify neurons as auditory, we compared the neuronal firing rates within a pre-stimulus window (-25 to 5 ms) to the firing rates during the stimulus or expected stimulus window (5 to 120 ms) using the Wilcoxon signed-rank test (signrank function in MATLAB). Neurons were classified as auditory if they showed a significant response (p < 0.05) to at least one of the tones (7 conditions with Tone A and 7 with Tone B, excluding the conditions with 0% of either tone) or their omission (8 conditions) in any of the experimental conditions, for a total of 22 comparisons. To control for false discoveries due to multiple comparisons, we applied the Benjamini-Hochberg procedure to adjust the p-values and account for the increased risk of Type I errors (false positives) when conducting multiple tests.

### ORN classification

To define omission responsiveness, we applied a similar test as described above, but only considered the omission responses across the eight experimental conditions. Neurons were classified as omission-responsive if they exhibited a significant response to omissions in at least one of the conditions, with p-values adjusted using the Benjamini-Hochberg procedure (the definition is similar to Lao-Rodriguez et al [26] defined as omission-sensitive neurons but they used one condition).

#### Omission preference

Due to the high prevalence of tone selectivity among ORNs, we defined the tone that elicited the strongest omission response from each neuron as the O_P_ tone, and the other tone as the O_NP_ tone. The omission response strength was determined by comparing the firing rates during the omission window (5 to 120 ms) over the baseline between the two 95% conditions.

#### Buildup of response

To analyze the dynamics of the omission response throughout the trials, we focused on the four experimental conditions where the O_P_ tone was the standard (95%, 90%, 85%, and 75% O_P_ tone). We averaged the response across these four conditions for each ORN and repetition, resulting in a 2D matrix representing the averaged response. We then computed the mean and standard error across ORNs for each repetition. The mean response across ORNs was fitted with a logistic model using the nlinfit function in MATLAB. The goodness of fit was assessed using the R-squared value, which was calculated by comparing the residual sum of squares to the total sum of squares.

#### PEON classification

To investigate whether neurons encode the probability of the omission-preferred tone, we calculated the Spearman rank correlation between the probability of the O_P_ tone and the neuronal activity during omissions. A one-tailed test was used to determine if higher O_P_ tone probabilities were associated with increased neuronal activity. Neurons exhibiting a significant positive correlation (p < 0.05) were defined as probability encoding.

#### Laminar and areal distribution of neurons

To determine the laminar distribution of neurons, we adapted the methodology from Shiramatsu et al. [47]. We used current source density (CSD) analysis on local field potentials recorded with Neuropixels electrodes to identify cortical layer boundaries. The earliest clear current sink, typically corresponding to the border between layers 3 and 4, was used as a key marker. Individual neuron depths were estimated based on their distance from the probe tip, as determined by spike sorting. Neurons were then classified into supragranular (Layers 1-3), granular (Layer 4), and infragranular (Layers 5-6) layers. Cortical thickness and layer boundaries were set according to Shiramatsu et al[47]

To assess the statistical significance of the observed laminar distribution, we employed a two-sided bootstrap analysis. We compared our actual distribution to 300,000 randomly generated distributions, maintaining the total number of neurons in each layer. P-values were calculated based on how often the random samples showed more extreme counts than our observed data, with p < 0.05 considered statistically significant. For the analysis of neuron distribution across different auditory areas (A1, VAF, AAF), we used a similar bootstrap approach, randomizing the assignment of neurons to different areas while maintaining the total number of neurons in each area. This method allowed us to determine whether the observed distribution of ORNs and PEONs across cortical layers and auditory areas was significantly different from what would be expected by chance, providing insight into the functional organization of these neurons in the auditory cortex.

### Computational modeling

We constructed a circuit of leaky integrate-and-fire neurons following

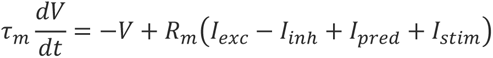

with membrane time constant 𝜏_𝑚_ = 30 ms and membrane resistance 𝑅_𝑚_ = 1 MΩ. When 𝑉 > 10 mV, neurons emitted a spike (𝑋𝑋 = 1) and were reset to 𝑉 = 0 mV for a refractory period of 2 ms. Synaptic currents 𝐼_𝑒𝑥𝑐_ and 𝐼_𝑖𝑛ℎ_ were modeled with instantaneous rise and exponential decay,

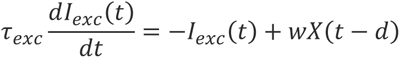

and equivalently for 𝐼_𝑖𝑛ℎ_, with time constants 𝜏_𝑒𝑥𝑐_ = 5 ms and 𝜏_𝑖𝑛ℎ_ = 20 ms. Spikes from presynaptic neurons 𝑋𝑋 ∈ {0,1} were delivered with a synapse-specific delay 𝑑 and weight 𝑤 as detailed in Supplemental Table S1 within each stream module. Lateral connections between streams 𝑠_1_ and 𝑠2 delivered spikes with delay d = 5|𝑠_2_ – 𝑠_1_ | ms and weight 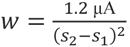. Sensory stimulation was presented through the input neuron I by setting its 𝐼_𝑠𝑡𝑖𝑚_ to a constant 𝑆_𝑠𝑡𝑟𝑒𝑛𝑔𝑡ℎ = 0.2 μA (target stream) or 𝑆_𝑠𝑡𝑟𝑒𝑛𝑔𝑡ℎ = 0.1 μA (adjacent streams) for 20 ms. Similarly, prediction signals were fed to the P neuron by setting 𝐼_𝑝𝑟𝑒𝑑_ = P_strength ∗ 0.2 μA with identical timing. *P_strength* was initialized to 0 and updated by spikes in prediction neurons as follows:

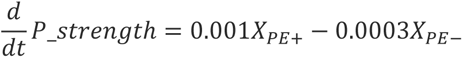

and clipped to ensure 0 ≤ 𝑃_𝑠𝑡𝑟𝑒𝑛𝑔𝑡ℎ ≤ 1 at all times. Both 𝐼_𝑠𝑡𝑖𝑚_ and 𝐼_𝑝𝑟𝑒𝑑_ were raised to their respective values for 20 ms at the start of each trial, then reset to 0 μA for the remaining 130 ms.

The model was written in Brian2 [50], which allowed calculating exact solutions for the internal neurons of the circuit (PE+, PE–, I+, I–). The remaining neurons (P and I), synapses and *P_strength* were integrated with a time step of 0.1 ms by forward Euler integration. Simulations were run over 500 trials.

## Data Availability

The datasets generated and analyzed during the current study are available from the corresponding author upon reasonable request during the manuscript submission and review process. Upon publication, these datasets will be accessible to readers in a public repository, with a DOI to be provided.

## Code Availability

The custom code and algorithms used in this study are available from the corresponding author upon reasonable request during the manuscript submission and review process. Upon publication, these codes will be accessible to readers in a public repository, with a DOI to be provided.

## Acknowledgments

We thank the International Research Center for Neurointelligence at the University of Tokyo for support and promotion of collaboration. This work was supported by World Premier International Research Center Initiative (WPI), MEXT, Japan, JSPS KAKENHI ( 23K14298, 23H03465, 23H04336, 23H03023, 24H01544), AMED (JP23dm0307009), JST (JPMJMS2296, JPMJPR22S8), the Asahi Glass Foundation, and the Secom Science and Technology Foundation.

## Author Contributions

Z.C.C. and A.Y. conceptualized the study. A.Y. and T.S. conducted the experiment, and A.Y. analyzed data. Z.C.C. developed the computational model, and F.B.K. helped with the model implementation. K.O. performed the free energy analysis. Z.C.C. and H.T. supervised the project. Z.C.C., H.T., A.Y., and T.S. acquired funding. A.Y., Z.C.C., and K.O wrote the first manuscript draft. H.T., T.S., and F.B.K. revised the manuscript. All authors contributed to the final version of the manuscript.

## Declaration of interests

The authors declare no competing interests.

Declaration of generative AI and AI-assisted technologies in the writing process During the preparation of this work, the authors used Claude.ai to improve readability. After using this tool, the authors reviewed and edited the content as needed and take full responsibility for the content of the published article.

## Supplemental Information

### Simplified Free Energy Principle

First, we assumed that the amplitudes of the sensory input (auditory tone) at the i-th frequency (ui) are generated by a single-hierarchy static linear generative model:

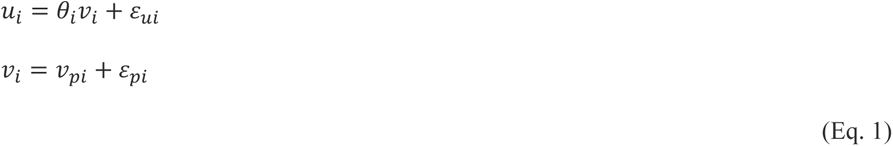

where v_i_ is a hidden variable to generate auditory tones at the i-th frequency, θ_i_ is a linear gain between vii and ui, and ε_ui_ and ε_pi_ are Gaussian noise. We assumed that the linear gain θ_i_, the prior expectation of the hidden variables 𝑣_𝑝𝑖_, and the covariance matrices Σ_𝑢_, Σ_𝑝_ are already adapted to the natural environment and stable over time, at least on the time scale of the electrophysiological experiments.

We assumed only one hierarchy in the predictive coding network because the generative model also has only one hierarchy. The estimate of the hidden variable at the i-th frequency (𝑣_𝑖_) is denoted as 𝜙_𝑖_.

According to Friston[51], hidden variables ϕI can be estimated by:

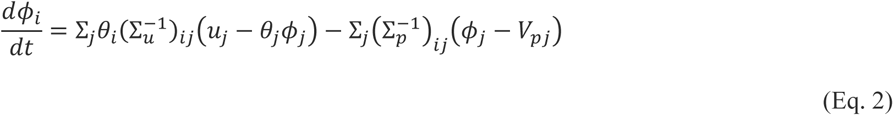

We defined signal, prediction, bias, and prediction error as follows:

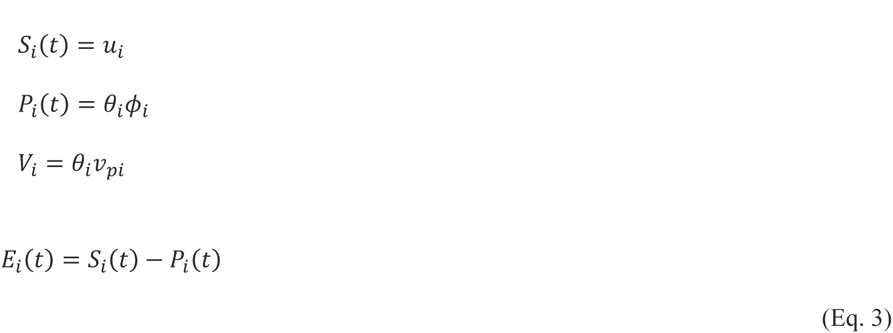

Si(t) is the sensory (auditory tone) amplitude at the i-th frequency,9 and Pi(t)i is the prediction of the tone amplitude at the i-th frequency. Vi is the bias due to the prior expectation of the tone amplitude at the i-th frequency. Ei(t) is the prediction error. We assumed that Si(t) is always positive or zero, because it represents the amplitude of the tone. The prediction error can be positive or negative and we separated the positive and negative prediction errors as follows:

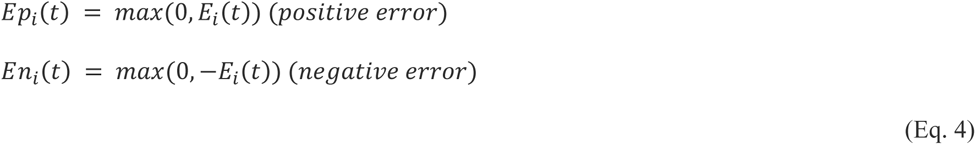

Then, Eq. 2 can be rewritten as follows:

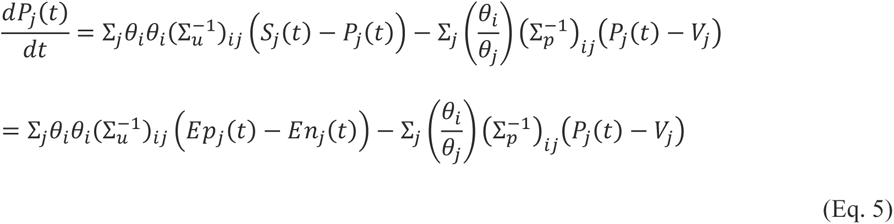

We assumed the correlation structures of ε_ui_ and ε_pi_ as follows:

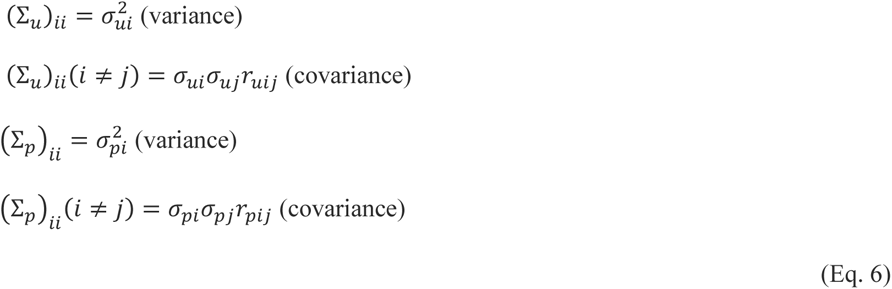

where ruij is a correlation coefficient between i-th frequency and j-th frequency.

When we considered only two tone frequencies, similar to the electrophysiological experiments, we can obtain the precision matrices (Σ_u_-1)_ij_ and (Σ_p_-1)_ij_ , that are the inverse of the covariance matrices as follows:

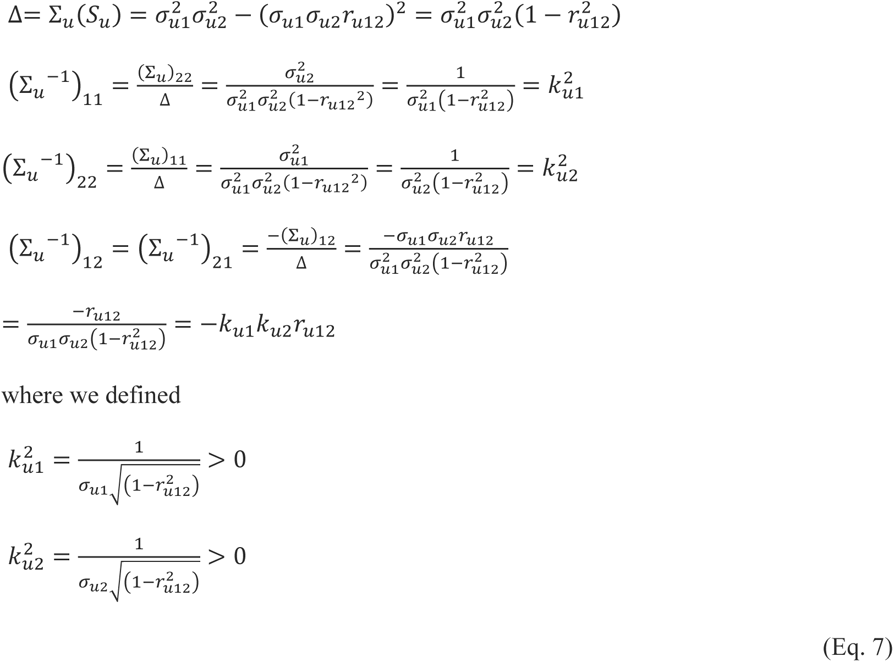

Similarly,

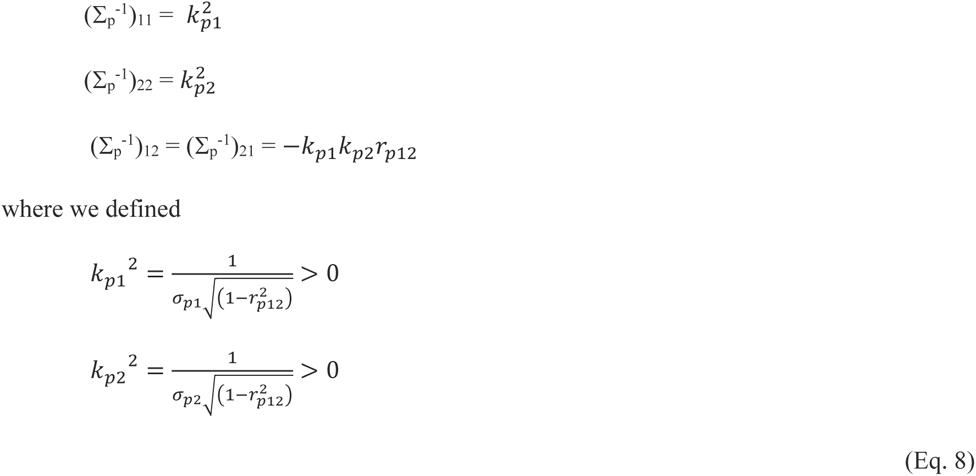

Then, Eq. 5 can be rewritten as follows:

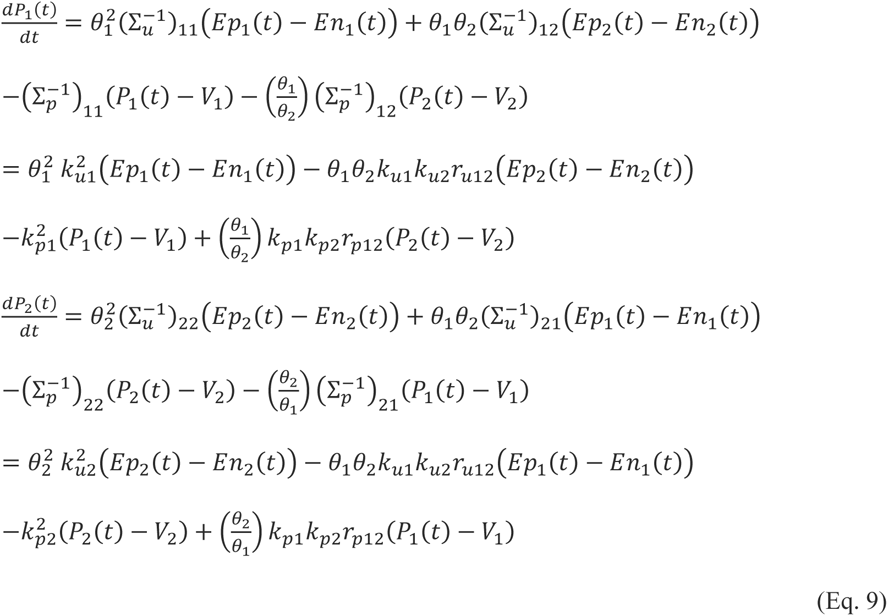

Further, we assumed that the noises of the hidden variables (ε_pi_) are independent over different frequencies:

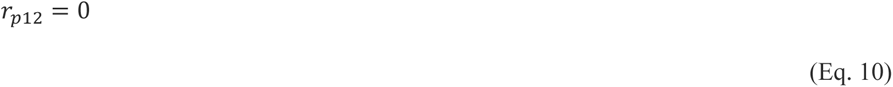

Then, we finally obtained the prediction update rule consisting of three terms:

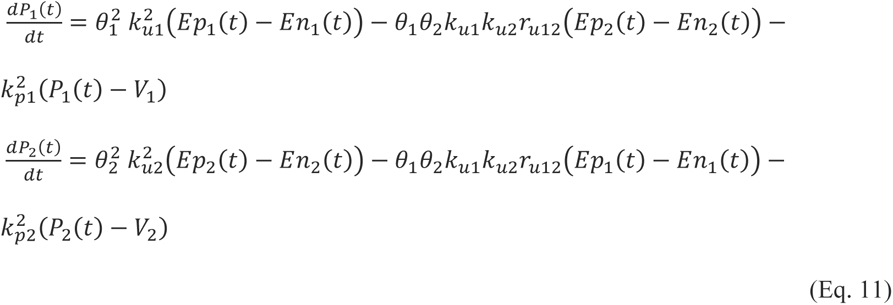

The first term, θ_1_^2^ k_u1_^2^ (Ep_1_(t) - En_1_(t)), represents the update by the positive and negative prediction errors in the same stream. The second term, – θ_1_ θ_2_ k_u1_ k_u2_ r_u12_ (Ep_2_(t) - En_2_(t)), represents the lateral interaction of positive and negative prediction errors across steams of the 1^st^ and 2^nd^ frequencies. The third term, – k_p1_^2^ (P_1_(t) –V_1_), represents the temporal decay of the prediction towards the prior expectation.

Thus, the lateral interaction of positive and negative errors across different streams is a natural outcome derived directly from the free energy principle. The sign of the lateral interaction depends on the correlation between the noise of the sensory input (ε_ui_) at the 1^st^ and 2^nd^ frequencies. In general, the amplitudes of tones at two different frequencies are positively correlated in the natural environment [52], the lateral interaction becomes suppressive.

---

**Figure S1.**
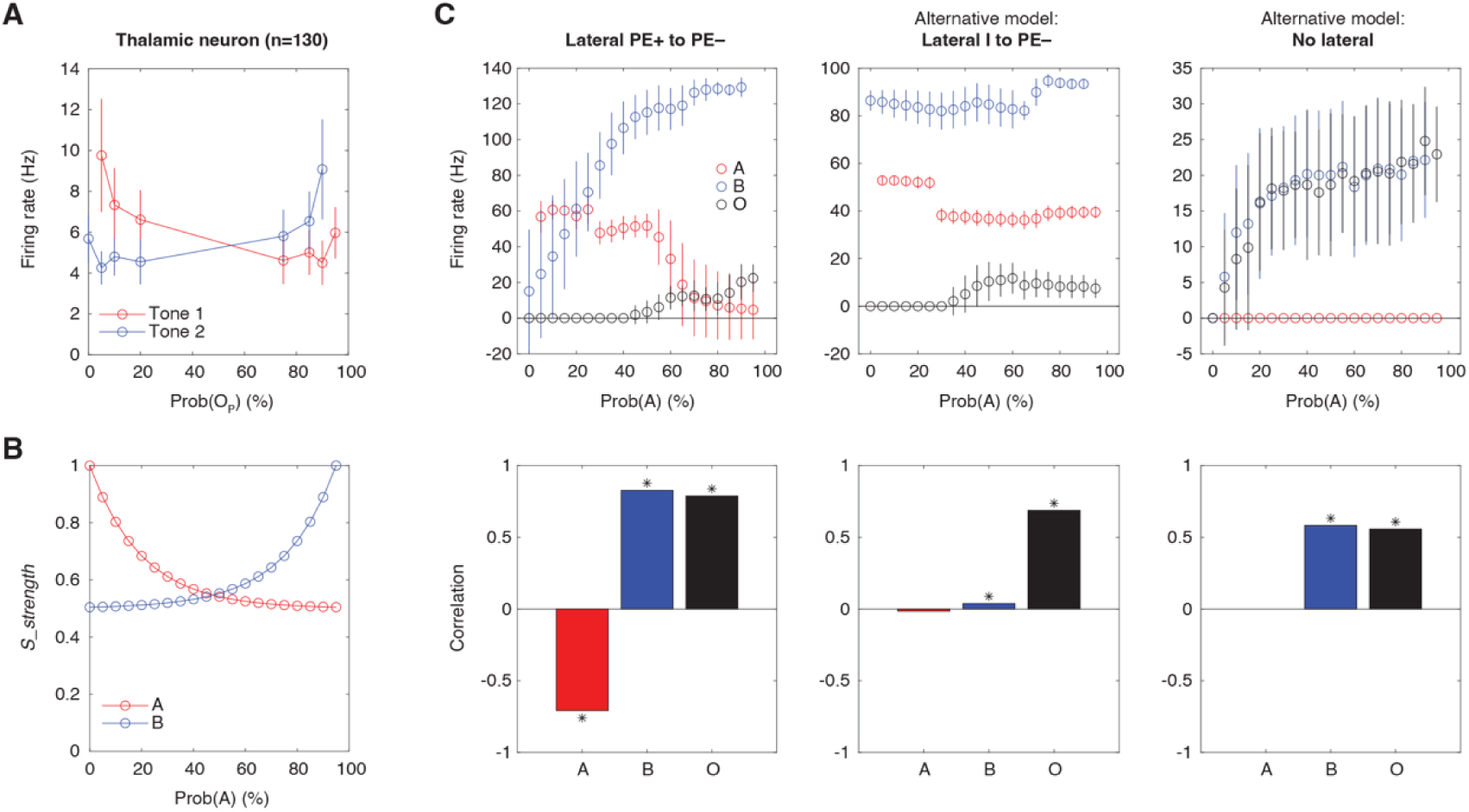
Model comparisons with adapted sensory signals. (**A**) Firing rates of thalamic neurons (n=130) for Tone 1 (red) and Tone 2 (blue) across varying probabilities of the preferred tone (Prob(OP)). Error bars indicate standard error of the mean (SEM). (**B**) To mimic the thalamic sensory inputs shown in the panel A, the strength of sensory signals (*S_strength*) for Tones A (red) and B (blue) was modelled as exponential decay functions. (**C**) Comparison of firing rates in different model configurations: Lateral PE+ to PE– (left), Lateral I to PE– (middle), and No lateral (right). Firing rates for Tones A (red), B (blue), and omissions (O, black) are plotted against Prob(A) in the top panels. Bottom panels display correlation coefficients between firing rates and probabilities for Tones A, B, and omissions, with significant correlations marked by asterisks. Error bars indicate SEM.

**Table S1.** Synaptic delays and weights within each stream module.

| Connection | Delay | Weight |
| --- | --- | --- |
| I to I− | 10 ms | 0.5 μA |
| I to PE+ | 20 ms | 0.5 μA |
| P to PE− | 20 ms | 0.5 μA |
| P to I+ | 10 ms | 0.5 μA |
| I− to PE− | 5 ms | 0.2 μA |
| I+ to PE+ | 5 ms | 0.2 μA |

## References

1. Friston K. The free-energy principle: a unified brain theory? Nature Reviews Neuroscience 2010 11:2. 2010;11: 127–138. doi:10.1038/nrn2787

2. Rao RPN, Ballard DH. Predictive coding in the visual cortex: a functional interpretation of some extra-classical receptive-field effects. Nature Neuroscience 1999 2:1. 1999;2: 79–87. doi:10.1038/4580

3. Friston K. A theory of cortical responses. Philos Trans R Soc Lond B Biol Sci. 2005;360: 815–836. doi:10.1098/RSTB.2005.1622

4. Bastos AM, Usrey WM, Adams RA, Mangun GR, Fries P, Friston KJ. Canonical microcircuits for predictive coding. Neuron. 2012;76: 695–711. doi:10.1016/j.neuron.2012.10.038

5. Clark A. Whatever next? Predictive brains, situated agents, and the future of cognitive science. Behavioral and Brain Sciences. 2013;36: 181–204. doi:10.1017/S0140525X12000477

6. Keller GB, Mrsic-Flogel TD. Predictive Processing: A Canonical Cortical Computation. Neuron. 2018;100: 424–435. doi:10.1016/J.NEURON.2018.10.003

7. Fiser A, Mahringer D, Oyibo HK, Petersen A V., Leinweber M, Keller GB. Experience-dependent spatial expectations in mouse visual cortex. Nat Neurosci. 2016;19: 1658– 1664. doi:10.1038/NN.4385

8. O’toole SM, Oyibo HK, Keller GB. PREDICTION ERROR NEURONS IN MOUSE CORTEX ARE MOLECULARLY TARGETABLE CELL TYPES. [cited 13 Mar 2024]. doi:10.1101/2022.07.20.500837

9. Leinweber M, Ward DR, Sobczak JM, Attinger A, Keller GB. A Sensorimotor Circuit in Mouse Cortex for Visual Flow Predictions. Neuron. 2017;95: 1420–1432.e5. doi:10.1016/J.NEURON.2017.08.036

10. Carbajal G V., Malmierca MS. The Neuronal Basis of Predictive Coding Along the Auditory Pathway: From the Subcortical Roots to Cortical Deviance Detection. Trends Hear. 2018;22. doi:10.1177/2331216518784822

11. Ulanovsky N, Las L, Nelken I. Processing of low-probability sounds by cortical neurons. Nat Neurosci. 2003;6: 391–398. doi:10.1038/NN1032

12. Yaron A, Hershenhoren I, Nelken I. Sensitivity to Complex Statistical Regularities in Rat Auditory Cortex. Neuron. 2012;76: 603–615. doi:10.1016/J.NEURON.2012.08.025

13. Chao ZC, Takaura K, Wang L, Fujii N, Dehaene S. Large-Scale Cortical Networks for Hierarchical Prediction and Prediction Error in the Primate Brain. Neuron. 2018;100: 1252–1266.e3. doi:10.1016/J.NEURON.2018.10.004

14. Garrido MI, Kilner JM, Stephan KE, Friston KJ. The mismatch negativity: a review of underlying mechanisms. Clin Neurophysiol. 2009;120: 453–463. doi:10.1016/J.CLINPH.2008.11.029

15. Shiramatsu TI, Kanzaki R, Takahashi H. Cortical Mapping of Mismatch Negativity with Deviance Detection Property in Rat. PLoS One. 2013;8: e82663. doi:10.1371/JOURNAL.PONE.0082663

16. Taaseh N, Yaron A, Nelken I. Stimulus-specific adaptation and deviance detection in the rat auditory cortex. PLoS One. 2011;6. doi:10.1371/JOURNAL.PONE.0023369

17. Wacongne C, Changeux JP, Dehaene S. A neuronal model of predictive coding accounting for the mismatch negativity. J Neurosci. 2012;32: 3665–3678. doi:10.1523/JNEUROSCI.5003-11.2012

18. Rubin J, Ulanovsky N, Nelken I, Tishby N. The Representation of Prediction Error in Auditory Cortex. Theunissen FE, editor. PLoS Comput Biol. 2016;12: e1005058. doi:10.1371/journal.pcbi.1005058

19. Bendixen A, Schröger E, Winkler I. I heard that coming: Event-related potential evidence for stimulus-driven prediction in the auditory system. Journal of Neuroscience. 2009;29: 8447–8451. doi:10.1523/JNEUROSCI.1493-09.2009

20. Chennu S, Noreika V, Gueorguiev D, Shtyrov Y, Bekinschtein TA, Henson R. Silent expectations: Dynamic causal modeling of cortical prediction and attention to sounds that weren’t. Journal of Neuroscience. 2016;36: 8305–8316. doi:10.1523/JNEUROSCI.1125-16.2016

21. Hughes HC, Darcey TM, Barkan HI, Williamson PD, Roberts DW, Aslin CH. Responses of human auditory association cortex to the omission of an expected acoustic event. Neuroimage. 2001;13: 1073–1089. doi:10.1006/nimg.2001.0766

22. Raij T, Mcevoy L, Makela JP, Harïbrain R, Harï H. Human auditory cortex is activated by omissions of auditory stimuli. Brain Res. 1997.

23. Yabe H, Tervaniemi M, Reinikainen K, Näätänen R. Introduction Temporal window of integration revealed by MMN to sound omission. 1997. Available: http://journals.lww.com/neuroreport

24. Heilbron M, Chait M. Great Expectations: Is there Evidence for Predictive Coding in Auditory Cortex? Neuroscience. Elsevier Ltd; 2018. pp. 54–73. doi:10.1016/j.neuroscience.2017.07.061

25. Awwad B, Jankowski MM, Polterovich A, Bashari S, Nelken I. Extensive representation of sensory deviance in the responses to auditory gaps in unanesthetized rats. Current Biology. 2023;33: 3024–3030.e3. doi:10.1016/j.cub.2023.06.013

26. Lao-Rodríguez AB, Przewrocki K, Pérez-González D, Alishbayli A, Yilmaz E, Malmierca MS, et al. Neuronal responses to omitted tones in the auditory brain: A neuronal correlate for predictive coding. 2023. Available: https://www.science.org

27. Auksztulewicz R, Rajendran VG, Peng F, Schnupp JWH, Harper NS. Omission responses in local field potentials in rat auditory cortex. BMC Biol. 2023;21. doi:10.1186/s12915-023-01592-4

28. Takahashi H, Nakao M, Kaga K. Interfield differences in intensity and frequency representation of evoked potentials in rat auditory cortex. Hear Res. 2005;210: 9–23. doi:10.1016/J.HEARES.2005.05.014

29. Takahashi H, Nakao M, Kaga K. Distributed representation of sound intensity in the rat auditory cortex. Neuroreport. 2004;15: 2061–2065. doi:10.1097/00001756-200409150-00013

30. Shiramatsu TI, Noda T, Akutsu K, Takahash H. Tonotopic and field-specific representation of long-lasting sustained activity in rat auditory cortex. Front Neural Circuits. 2016;10: 201972. doi:10.3389/FNCIR.2016.00059/BIBTEX

31. Friston K, Kilner J, Harrison L. A free energy principle for the brain. J Physiol Paris. 2006;100: 70–87. doi:10.1016/j.jphysparis.2006.10.001

32. Bendixen A, SanMiguel I, Schröger E. Early electrophysiological indicators for predictive processing in audition: A review. International Journal of Psychophysiology. 2012. pp. 120–131. doi:10.1016/j.ijpsycho.2011.08.003

33. Blakemore C, Tobin EA. Lateral inhibition between orientation detectors in the cat’s visual cortex. Exp Brain Res. 1972;15: 439–440. doi:10.1007/BF00234129

34. Metherate R, Kaur S, Kawai H, Lazar R, Liang K, Rose HJ. Spectral integration in auditory cortex: mechanisms and modulation. Hear Res. 2005;206: 146–158. doi:10.1016/J.HEARES.2005.01.014

35. Read HL, Winer JA, Schreiner CE. Functional architecture of auditory cortex. Curr Opin Neurobiol. 2002;12: 433–440. doi:10.1016/S0959-4388(02)00342-2

36. Schreiner CE, Read HL, Sutter ML. MODULAR ORGANIZATION OF FREQUENCY INTEGRATION IN PRIMARY AUDITORY CORTEX. Annu Rev Neurosci. 2000;23: 501–529.

37. Ojima H, Honda C, Cortex EJ-C, 1991 undefined. Patterns of axon collateralization of identified supragranular pyramidal neurons in the cat auditory cortex. academic.oup.comH Ojima, CN Honda, E JonesCerebral Cortex, 1991•academic.oup.com. [cited 19 Jul 2024]. Available: https://academic.oup.com/cercor/article-abstract/1/1/80/408924

38. Budinger E, Scheich H. Anatomical connections suitable for the direct processing of neuronal information of different modalities via the rodent primary auditory cortex. Hear Res. 2009;258: 16–27. doi:10.1016/J.HEARES.2009.04.021

39. Kaur S, Lazar R, Metheate R. Intracortical pathways determine breadth of subthreshold frequency receptive fields in primary auditory cortex. J Neurophysiol. 2004;91: 2551– 2567. doi:10.1152/JN.01121.2003

40. Happel MFK, Jeschke M, Ohl FW. Spectral Integration in Primary Auditory Cortex Attributable to Temporally Precise Convergence of Thalamocortical and Intracortical Input. Journal of Neuroscience. 2010;30: 11114–11127. doi:10.1523/JNEUROSCI.0689-10.2010

41. Levy RB, Reyes AD. Spatial Profile of Excitatory and Inhibitory Synaptic Connectivity in Mouse Primary Auditory Cortex. Journal of Neuroscience. 2012;32: 5609–5619. doi:10.1523/JNEUROSCI.5158-11.2012

42. Wertz A, Trenholm S, Yonehara K, Hillier D, Raics Z, Leinweber M, et al. PRESYNAPTIC NETWORKS. Single-cell-initiated monosynaptic tracing reveals layer-specific cortical network modules. Science. 2015;349: 70–74. doi:10.1126/SCIENCE.AAB1687

43. Stettler DD, Das A, Bennett J, Gilbert CD. Lateral connectivity and contextual interactions in macaque primary visual cortex. Neuron. 2002;36: 739–750. doi:10.1016/S0896-6273(02)01029-2

44. Narayanan RT, Udvary D, Oberlaender M. Cell type-specific structural organization of the six layers in rat barrel cortex. Front Neuroanat. 2017;11. doi:10.3389/FNANA.2017.00091/PDF

45. Chang M, Kanold PO. Development of Auditory Cortex Circuits. JARO: Journal of the Association for Research in Otolaryngology. 2021;22: 237. doi:10.1007/S10162-021-00794-3

46. Harpaz M, Jankowski MM, Khouri L, Nelken I. Emergence of abstract sound representations in the ascending auditory system. Prog Neurobiol. 2021;202. doi:10.1016/j.pneurobio.2021.102049

47. Shiramatsu TI, Takahashi K, Noda T, Kanzaki R, Nakahara H, Takahashi H. Microelectrode mapping of tonotopic, laminar, and field-specific organization of thalamo-cortical pathway in rat. Neuroscience. 2016;332: 38–52. doi:10.1016/J.NEUROSCIENCE.2016.06.024

48. Jun JJ, Steinmetz NA, Siegle JH, Denman DJ, Bauza M, Barbarits B, et al. Fully integrated silicon probes for high-density recording of neural activity. Nature 2017 551:7679. 2017;551: 232–236. doi:10.1038/nature24636

49. Pachitariu M, Steinmetz N, Kadir S, Carandini M, D. HK. Kilosort: realtime spike-sorting for extracellular electrophysiology with hundreds of channels. bioRxiv. 2016; 061481. doi:10.1101/061481

50. Stimberg M, Brette R, Goodman DFM. Brian 2, an intuitive and efficient neural simulator. Elife. 2019;8. doi:10.7554/ELIFE.47314

51. Friston K. A theory of cortical responses. Philosophical Transactions of the Royal Society B: Biological Sciences. 2005;360: 815–836. doi:10.1098/RSTB.2005.1622

52. Nelken I, Rotman Y, Yosef OB. Responses of auditory-cortex neurons to structural features of natural sounds. Nature. 1999;397: 154–157. doi:10.1038/16456

